# CEREBRUM-7T: Fast and Fully-volumetric Brain Segmentation of 7 Tesla MR Volumes

**DOI:** 10.1101/2020.07.07.191536

**Authors:** Michele Svanera, Sergio Benini, Dennis Bontempi, Lars Muckli

## Abstract

Ultra high-field MRI enables sub-millimetre resolution imaging of the human brain, allowing for the resolution of functional circuits at the meso-scale of cortical layers. An essential step in many functional and structural neuroimaging studies is segmentation, the operation of partitioning the MR brain images to delineate anatomical structures. Despite recent efforts in brain imaging analysis, the literature lacks of accurate and fast methods for segmenting 7 Tesla (7T) brain MRI. We here present `CEREBRUM-7T`, an optimised end-to-end Convolutional Neural Network (CNN) architecture, that allows for the segmentation of a whole 7T T1_w_ MRI brain volume at once, thus overcoming the drawbacks of partitioning the volume into 2D or 3D tiles. Training is performed in a weakly supervised fashion, exploiting labelling with errors obtained with automatic state-of-the-art methods. The trained model is able to produce accurate multi-structure segmentation masks on six different classes in only a few seconds. In the experimental part, a combination of objective numerical evaluations and subjective analysis carried out by experienced neuroimaging users, confirms that the proposed solution outperforms the training labels it was trained on in segmentation accuracy, and is suitable for neuroimaging studies, such as layer fMRI studies. Taking advantage of a fine-tuning operation on a reduced set of volumes, we also show how it is possible to efficiently and effectively apply `CEREBRUM-7T` to data from different sites. Furthermore, to allow replicability and encourage extensions, we release the code, 7T data (142 volumes), and other materials, including the training labels and the Turing test.

## 1 Introduction

Magnetic Resonance Imaging (MRI) is a method for non-invasively measuring brain anatomy and function, widespread in research and clinical environments. Most MRI scanners in clinical practice have a field strength of 3 Tesla or less. However, ultra-high field magnetic resonance machines, with a field strength of 7T, are now CE labeled and FDA approved, and becoming more common in research and clinical settings. The introduction of 7T scanners, together with improvements in acquisition methods, has led to substantial advances in both signal and contrast to noise ratio, and increased the imaging resolution to a sub-millimetre level (Duyn, 2012). This technology enables imaging of different cortical depths and columns in order to investigate functional circuits of human cortex with high spatial specificity, and the visualisation of structures with an unprecedented definition.

However, the capability of using these innovative systems also poses new challenges, including standardisation across 7T sites, despite recent efforts show improved consistency by synchronising sequence protocols across centres; an example is the UK7T Network (Clarke et al., 2020). Furthermore, standardised brain atlases and intensity-based pipelines specific for the analysis of 7T data will help to minimise variability in analyses across sites (Botvinik-Nezer et al., 2020). Other problems in ultra-high field MR imaging concern the susceptibility artefacts that intervene during acquisition (Uğurbil et al., 2003), and the inhomogeneities in the magnetic field that lead to intensity variations across the image. These effects represent particular challenges for the analysis of structural data: for example, non-uniform voxel intensities within a given MRI can lead to low quality processing outcomes.

Image segmentation is an essential quantitative analysis step when assessing both healthy brain anatomy and patho-physiological conditions. The segmentation of brain structures is of paramount importance for monitoring anatomical variations during the development of neuro-degenerative processes, psychiatric disorders, and neurological diseases. Additionally, segmentation is an essential step in functional MRI (fMRI) studies, to isolate specific brain regions where to investigate patterns of brain activity. For example, in layer fMRI studies, being able to accurately segment inner and outer grey matter boundaries is of crucial importance.

### 1.1 Existing methods for 7T brain segmentation

Popular fully-automatic segmentation tools available for 3T data - such as FreeSurfer (Fischl, 2012) - which usually apply atlas-based or multi-atlas-based segmentation strategies, are not maximally effective on 7T volumes, due to increased data complexity, voxel intensity inhomogeneity, and different intensity distributions across sites. For example, as we will show later on, the last version of FreeSurfer (v7), which has been improved for UHF data, is able to segment very well multiple areas (e.g., GM/WM boundary), but the inhomogeneity of the scan affects its ability to correctly segment every region of the brain.

As a consequence, manual segmentation protocols (Wenger et al., 2014; Berron et al., 2017), although time consuming (Zhan et al., 2018; Koizumi et al., 2019), are still a common practise for 7T data. To partially reduce the laboursome process of manual segmentation, the solution proposed by Gulban et al. (2018) combines manual and semi-automatic segmentation, by adopting a multi-dimensional transfer function to single out non-brain tissue voxels in 7T MRI data of nine volunteers. Other semi-automated methods developed in the past for generic MRI data, such as `ITK-SNAP` (Yushkevich et al., 2006), have been adapted by (Archila-Meléndez et al., 2018) for tackling also ultra-high field brain imaging.

Often, given the lack of harmonised neuroimaging analysis protocols, multiple 7T sites created in-house pipelines to perform MRI segmentation on site-data specifically. Fracasso et al. (2016) developed a custom workflow (used for example in Bergmann et al., 2019) which we also adopt for labelling gray and white matter (GM and WM). The placement of the GM/CSF (cerebrospinal fluid) boundary is based on the location of the 75% quantile of the variability of T1_w_ partial volume estimates across cortical depth, while GM/WM boundary from a combination of `AFNI-3dSeg` (Cox, 1996) and a clustering procedure (see Section 2.2 for more details).

As an attempt to develop a site-independent approach, Bazin et al. (2014) presented a computational framework for whole brain segmentation at 7T, specifically optimised for MP2RAGE sequences. The authors develop a rich atlas of brain structures, on which they combine a statistical and a geometrical model. The method, which includes a non-trivial preprocessing chain for skull stripping and dura estimation, achieves whole brain segmentation and cortical extraction, all within a computation time below six hours. Despite these efforts, most existing solutions, including Bazin et al. (2014) and Nighres by Huntenburg et al. (2018), still generate a variety of segmentation errors that needs to be manually addressed, as reported in Gulban et al. (2018).

### 1.2 Deep learning methods for MRI segmentation

In recent years, the advanced classification and segmentation capabilities ensured by deep learning (DL) methods have impacted several medical imaging domains (Litjens et al., 2017; Hamidinekoo et al., 2018). Various segmentation algorithms, which exploit the generalization capabilities of DL and Convolutional Neural Networks (CNN) on unseen data, made possible a drastic improvement in the performance with respect to other traditional, mostly atlas-based, segmentation tools.

To the best of our knowledge, no DL architectures have been directly applied on 7T data for segmentation purposes yet. The only attempt made by Bahrami et al. (2016) to use CNN in this field, aimed at reconstructing 7T-like images from 3T MRI data. Specifically, from the 3T image intensities and the segmentation labels of 3T patches, the CNN learns a mapping function so as to generate the corresponding 7T-like image, with quality similar to the ground-truth 7T MRI.

Restricting the scope to 3T data only, recent DL-based methods such as QuickNat (Roy et al., 2019), MeshNet (Fedorov et al., 2017; McClure et al., 2019), NeuroNet (Rajchl et al., 2018), DeepNAT (Wachinger et al., 2018), and FastSurfer (Henschel et al., 2020) have been the most effective solutions among those which proposed to obtain a whole brain segmentation. However, a common trait of all the aforementioned methods is that none of them fully exploit the 3D spatial nature of MRI data, thus making segmentation accuracy sub-optimal. In fact, such solutions partition the brain into 3D sub-volumes (DeepNAT, MeshNet, and NeuroNet) or 2D patches (QuickNAT, FastSurfer), which are processed independently and only eventually reassembled; as recently shown in Reina et al. (2020), the use of “tiling” introduces small but relevant differences during inference that can negatively affect the overall quality of the segmentation result. For example in MRI segmentation, tiling entails a loss of global contextual information, such as the absolute and relative positions of different brain structures, which negatively impacts the segmentation outcome.

The DL model `CEREBRUM` (Bontempi et al., 2020) is the first attempt to fully exploit the 3D nature of MRI 3T data, taking advantage of both global and local spatial information. This 3D approach adopts an end-to-end encoding/decoding fully convolutional structure. Trained in a weakly supervised fashion with 900 whole brain volumes segmented with FreeSurfer v6 (Fischl, 2012), `CEREBRUM` learns to segment out-of-the-scanner^2^ brain volumes, with neither atlas-registration, pre-processing, nor filtering, in just ~5-10 seconds on a desktop GPU.

### 1.3 Aims and contributions

In this work we adapt and extend the framework presented in Bontempi et al. (2020) to handle UHF high-resolution data, delivering `CEREBRUM-7T`, the first DL fully automatic solution for brain MRI segmentation on out-of-the-scanner 7T data.

Similarly to its predecessor on 3T data Bontempi et al. (2020), `CEREBRUM-7T` acts in a fully 3D fashion on brain volumes, as shown in Figure 1. Despite the increase in the data size with respect to 3T scans, the volumetric processing of the whole brain is still possible by simplifying the model architecture with respect to other DL-based segmentation methods, and by distributing different part of the model on different GPUs. Such design choice enables to achieve a full brain segmentation in few seconds compared to several hours of other currently used non-DL methods for 7T data. Furthermore, exploiting the ability of DL methods to efficiently learn internal representations of data, brain segmentation happens with neither the support of ad-hoc preprocessing, nor the alignment to reference atlases.

**Figure 1.**
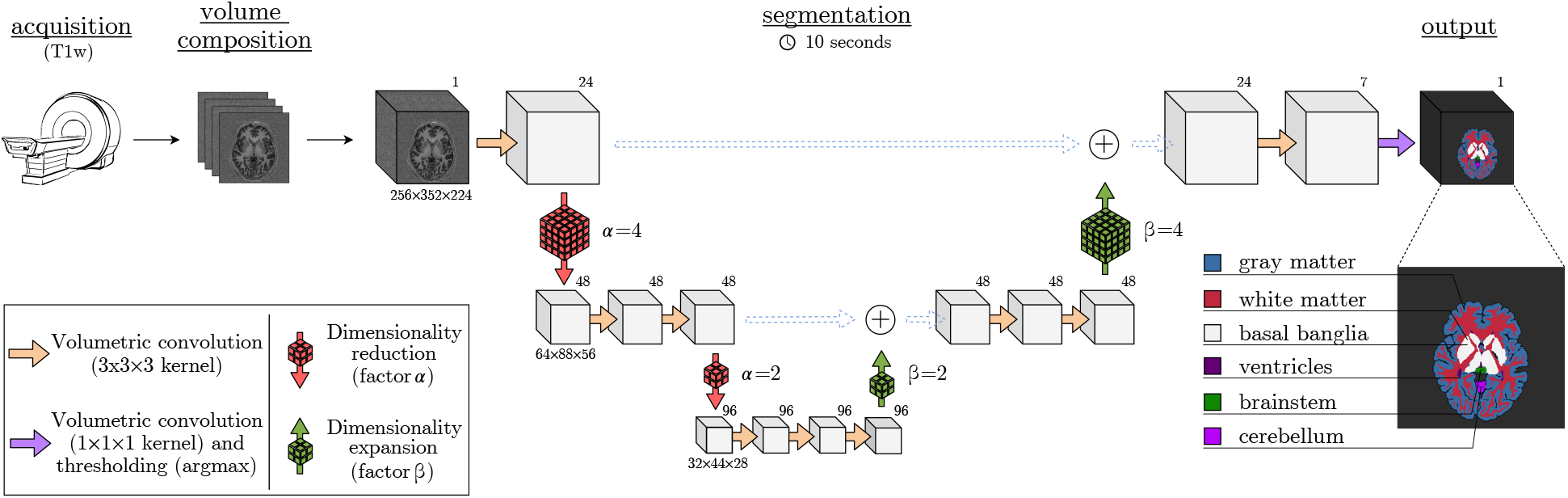
Method overview. The database is composed of 142 T1_w_ volumes (MP2RAGE at 7T, 0.63 *mm*^3^-iso) and an inaccurate ground-truth (iGT) is obtained using a combination of `AFNI - 3dSeg` (Cox, 1996) and methods as in Fracasso et al. (2016). Segmentation results are provided in only few seconds (~ 10 on a desktop CPU). Data dimensions are provided below each volume, while the numbers of filters are in the top-right corners.

The database we use is composed by 142 anatomical scans collected in Glasgow (UK), which currently constitutes the largest brain MRI 7T database for segmentation openly available. However, differently from Bontempi et al. (2020), where the dimension of the dataset was well enough for training (900 volumes), the scarcity of 7T volumes for training `CEREBRUM-7T` architecture, forced us to implement an ad-hoc data augmentation strategy.

The model is trained in a weakly-supervised fashion by exploiting a labelling with errors, which in the following we will refer to as *inaccurate* Ground Truth (iGT). Differently from the classical nomenclature for ground-truth (GT), we sharpen the name adding a clear indication that the training labels exhibit multiple errors. Such iGT is obtained by an in-house procedure which combines current state-of-the-art methods, and adopts the labelling strategy proposed by the MICCAI challenge (Mendrik et al., 2015): gray matter (GM), white matter (WM), cerebrospinal fluid (CSF), ventricles, cerebellum, brainstem, and basal ganglia. The iGT segmentation masks for GM/WM are obtained by combining geometric and clustering approaches (Cox, 1996; Fracasso et al., 2016), while other labels by using atlas-based techniques (Fischl, 2012) adequately processed, corrected, and adapted for 7T data. Although available anatomical labelling exhibits several errors, we show how the adopted learning procedure, exploiting “inaccurate supervision” (Zhou, 2017), ensures good generalisation capabilities and improved labelling performance with respect to the iGT it was trained on.

In the experimental part, `CEREBRUM-7T` is first quantitatively compared against the reference iGT, showing the beneficial effects of the data augmentation strategies. Results are presented by the same metrics used in the MICCAI MR Brain Segmentation challenge, i.e., the Dice Similarity Coefficient, the 95th Hausdorff Distance, and the Volumetric Similarity Coefficient (Mendrik et al., 2015). We then show that seven expert neuroscientists, involved in a Turing test (i.e., a blind survey, also referred to as *behavioural* in neuroscience) performed on three testing volumes which we manually segmented, consider `CEREBRUM-7T` segmentations better than the reference iGT used to train the model. Segmentation masks compared in the Turing test are also numerically examined using the above mentioned MICCAI metrics, on 2.7*M* annotated voxels, by comparing the results obtained by `CEREBRUM-7T`, FreeSurfer v6, FreeSurfer v7, the method in Fracasso et al. (2016), Nighres by Huntenburg et al. (2018), and the reference iGT used for training. Such experimental part, which is a combination of objective numerical evaluations and subjective analysis carried out by experienced neuroscientists, assesses the superiority of `CEREBRUM-7T` segmentation masks with respect to previous state-of-the-art methods.

With the objective of releasing a tool which is effective and efficient also on data from new sites, we eventually demonstrate the practical portability of the trained model, also on limited size data, by performing two fine-tuning experiments, one on AHEAD data Alkemade et al. (2020) (20 volumes), and the other on the Schneider et al. (2019) dataset (4 volumes).

As a last contribution, we make publicly available through the project website the set of 142 7T Glasgow scans^3^, the related iGT, all the code necessary to train and fine-tune `CEREBRUM-7T`, and to perform the Turing test.

## 2 Material and methods

### 2.1 Glasgow data: acquisition and split

The database consists of 142 out-of-the-scanner volumes obtained with a MP2RAGE sequence at 0.63 *mm*^3^ isotropic resolution, using a 7-Tesla MRI scanner with 32-channel head coil. All volumes were collected, as reconstructed DICOM images, at the Imaging Centre of Excellence (ICE) at the Queen Elizabeth University Hospital, Glasgow (UK). The columns of Figure 2 show some selected slices of the out-of-the-scanner T1_w_, the segmentation resulting from FreeSurfer v6, the one from Fracasso et al. (2016), the one from Huntenburg et al. (2018), the reference iGT, the `CEREBRUM-7T` mask, and the manual segmentation, respectively.

**Figure 2.**
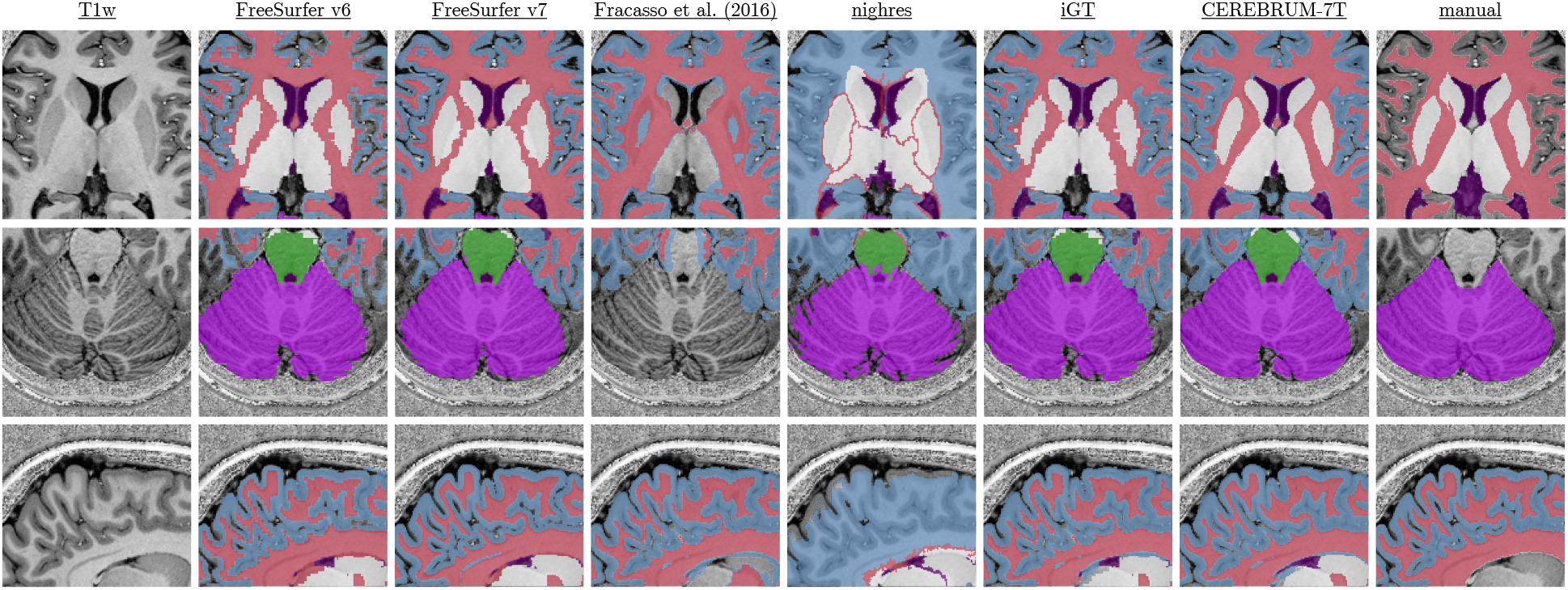
Visual examples of the dataset and results. Columns, from left to right, show: T1_w_ scan (left), FreeSurfer v6 and v7 segmentations, Fracasso et al. (2016), nighres, iGT (obtained as described in Section 2.2), `CEREBRUM-7T`, and manual segmentation. Coloured labels in are shown in overlay only when returned by the specific method.

Out of the total 142 volumes, 110 are used for training, 6 for validation, and 26 for testing (3 of which used in the Turing test). The only preprocessing applied is the neck cropping using the INV2 scan obtained during acquisition. Dataset details are shown in Table 1.

**Table 1.** Dataset details. MR Images acquired at the Imaging Centre of Excellence (ICE) at the Queen Elizabeth University Hospital, Glasgow (UK).

| Parameter | Value |
| --- | --- |
| Sequence used | T1<sub>w</sub> MP2RAGE |
| Field strenght | 7 Tesla |
| Voxel size | $0.63 \times 0.625 \times 0.625$ |
| Original volume sizes | $256 \times 360 \times 384$ |
| Training volume sizes | $256 \times 352 \times 224^{\dagger}$ |
| Training | 110 vol. (+1.1k vol. augmented offline) |
| Validation | 6 volumes |
| Testing | 26 volumes |
| Testing (Turing test) | 3 vol. × 8 areas |

<sup>†</sup> neck cropping

### 2.2 Generation of the inaccurate ground-truth

Similarly to most approaches employing DL frameworks for brain MRI segmentation (Roy et al., 2019; McClure et al., 2019; Fedorov et al., 2017; Rajchl et al., 2018; Bontempi et al., 2020), we also adopt an almost fully automatic procedure for labelling, since the prohibitive time cost required to produce a manual annotation on such large dataset. Such a decision is also driven by the consideration that, despite the use of an inaccurate ground truth (iGT), in already documented cases the trained models proved to perform the same (Rajchl et al., 2018), or even better (Bontempi et al., 2020; Roy et al., 2019), than the inaccurate ground-truth used for training.

Differently from Bontempi et al. (2020), we cannot use FreeSurfer (Fischl, 2012) as the only tool for segmenting all tissue and sub-cortical brain structures. As also hinted in Figure 2 (second column) the quality of WM and GM segmentation masks obtained with such tool is not acceptable for our purposes, even considering inaccurate supervision for learning.

An overview of the iGT generation process is shown in Figure 3. The pipeline accounts for two main branches: the upper one deals with WM and GM segmentation, while the lower one isolates other brain structures such as cerebellum, ventricles, brainstem, and basal ganglia. The two processing branches are combined afterwards, when a manual correction step is also carried out to reduce major errors.

**Figure 3.**
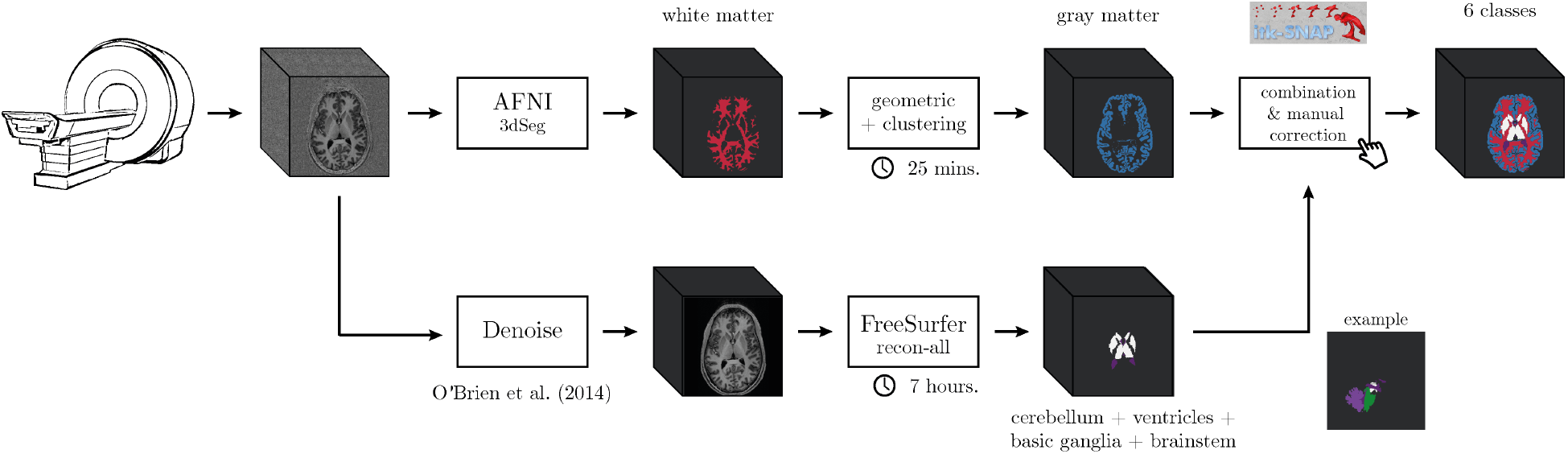
Processing pipeline used to generate the iGT starting from the reconstructed T1_w_.

In the upper branch the white matter (WM) mask is obtained using a combination of `AFNI - 3dSeg` (Cox, 1996) followed by geometric and clustering methods as in Fracasso et al. (2016). Specifically T1_w_ images are co-registered to an atlas (Desikan et al., 2006) and a brain mask is overlaid to the T1_w_ images to remove the cerebellum and subcortical structures. The T1_w_ images are then separated in six different parts along the posterior to anterior direction to improve intensity homogeneity. Each part is afterwards separately processed by the `3dSeg` function in `AFNI`, to isolate WM. The WM masks obtained from each part are summed together resulting in whole brain WM mask (see Fracasso et al. (2016) for further details).

The GM segmentation exploits such whole brain WM segmentation and an atlas co-registered to the T1_w_ images (Desikan et al., 2006). Next, a distance map from the WM/GM boundary to the pial surface is built computing the Euclidean distance of each voxel from the WM/GM border. Negative distances are assigned inside WM and positive distances are assigned from WM borders onward. Each region of interest (ROI) in the atlas by Desikan et al. (2006) is selected iteratively. For each ROI the coordinates are divided into four separate subparts using k-means clustering. For each subpart, voxels within −2*mm* and 7*mm* (Euclidean distance) from the WM/GM border are selected and their T1_w_ intensity stored for further analysis. For each cluster 10 bins between −2*mm* and 7*mm* are obtained – with each bin containing 10% of the data. For each of them, a partial volume estimate, defined as the standard deviation of T1_w_ intensity as well as the average Euclidean distance for the same bin, is computed. A linear model is then fit between the average Euclidean distance of each bin and the corresponding partial volume estimate. The slope of the linear model can be either positive or negative: if there is a positive slope, the 75% quantile of the standard deviation values is computed among the 10 bins; if, on the other hand, the slope is negative, the 25% quantile is computed. The Euclidean distance of the 25%/75% quantile corresponds to a drop or rise in T1_w_ variability and is considered as the transition between GM and CSF. To improve the obtained gray matter segmentation, the WM and GM masks are fed to the Cortical Reconstruction using Implicit Surface Evolution (CRUISE) algorithm in the Nighres software package (Huntenburg et al., 2018).

Despite the method ensures a robust result in segmenting GM/WM boundary, no cerebellum, ventricles, and basal ganglia areas are computed. To address such lack of GT structures, in the lower branch of the pipeline (Figure 3), we use FreeSurfer v6 (Fischl, 2012), which first requires to denoise the T1_w_ volume (O’Brien et al., 2014), to add the following labels: cerebellum, ventricles, brainstem, and basal ganglia. When necessary, a final step of manual correction is carried out to reduce major errors (especially on cerebellum classified as GM) using `ITK-SNAP` (Yushkevich et al., 2006).

The z-scoring procedure, applied to normalise the data before `CEREBRUM-7T` training, is obtained using the mean and standard deviation volumes computed on the entire Glasgow dataset (shown in Figure 4).

**Figure 4.**
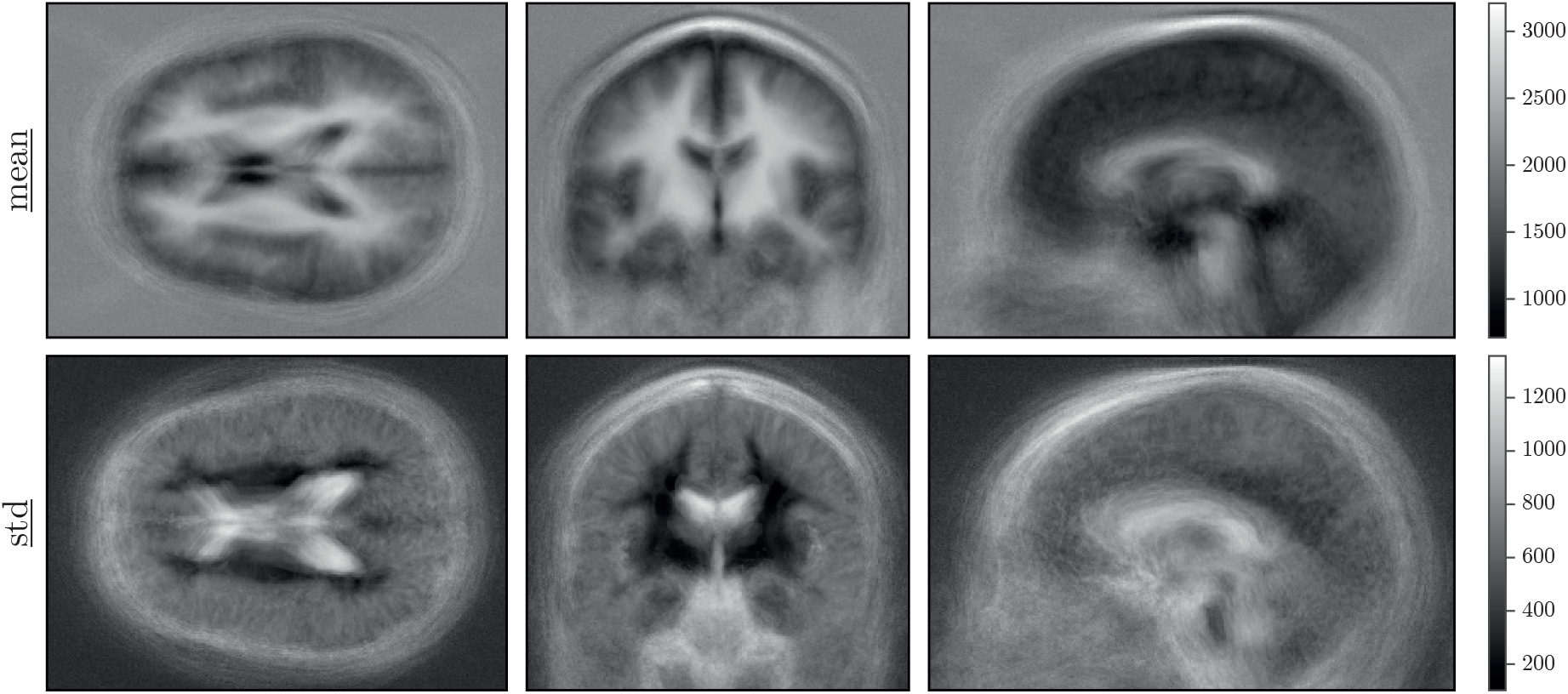
Mean and standard deviation volumes of the database used to z-score the data. Denoised mean/std volumes are found in Supplementary Material.

#### 2.2.1 Data augmentation

The data corpus used for our experiments is one of the biggest 7T brain MRI publicly and freely available datasets. Yet, given the complexity of the DL architecture, i.e., the number of learnable parameters, there are not enough training samples to deliver an off-the-shelf model. Therefore, we decide to adopt two customised data augmentation strategies: offline and online data augmentation, as shown in Figure 5.

**Figure 5.**
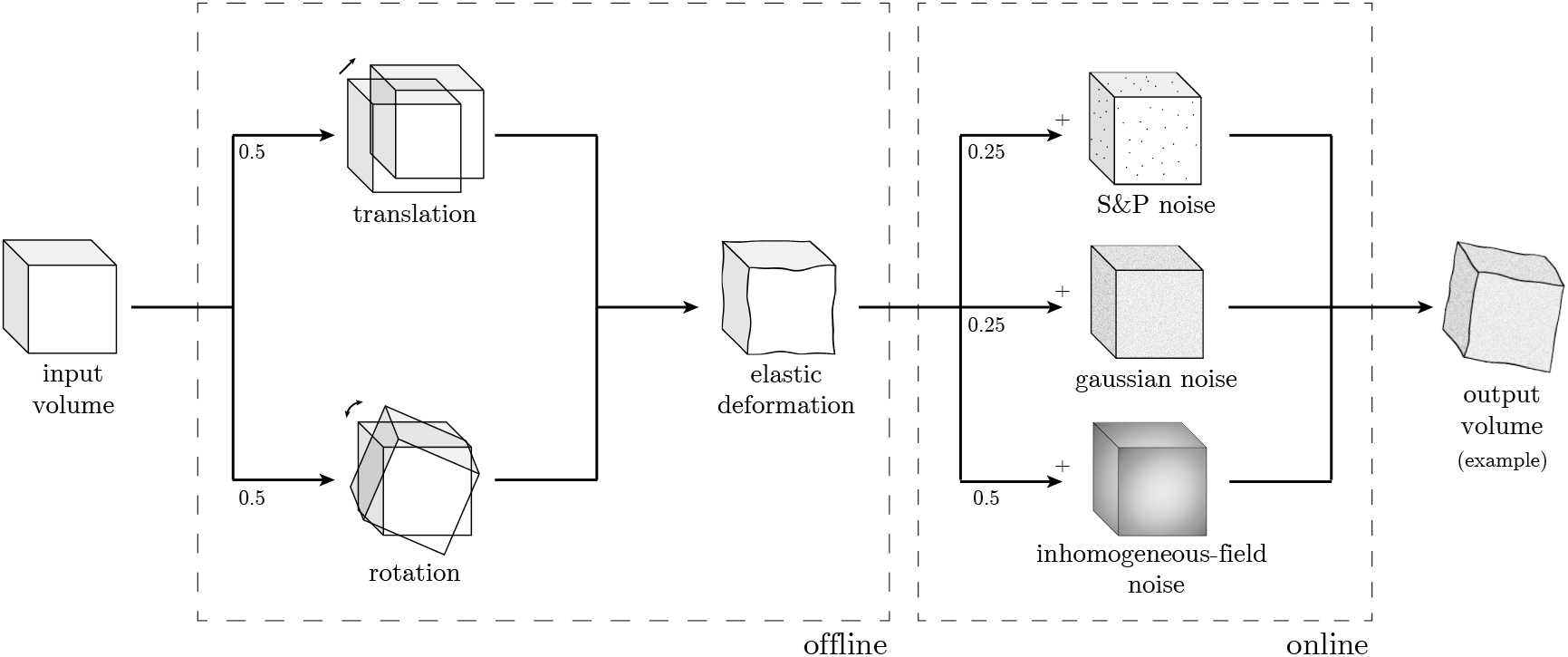
Data augmentation procedure. Offline (with respect to the training procedure), geometric augmentation is performed with rotation or translation (*p* = {0.5, 0.5}) and elastic deformation (Çiçek et al., 2016). Online, voxel intensity changes are applied: salt and pepper noise (*p* = 0.25), gaussian noise (*p* = 0.25), or inhomogeneous-field noise (*p* = 0.5). An example of data augmentation is shown in the output volume, where one rotation and one elastic deformation are followed by an additional gaussian noise.

Offline augmentation, too computationally demanding to be performed in training-time, consists in the application of small random shifts (*max*_*shift* = [10, 15, 10] voxels) or rotations (*max*_*rotation* = 5°, on all three axes), and elastic deformations. This ensures an augmentation factor of 10 of the training set.

Online data augmentation, performed during training, comprises variations on voxel intensities only: gaussian, salt & pepper, and inhomogeneous-field noise. In MRI, and especially in UHF, the inhomogeneity in the magnetic field produces an almost linear shift in the voxel intensity distributions for different areas in the 3D space (Sled et al., 1998). In other words: the same anatomical structure has different voxel intensities in different areas, for example GM in frontal and occipital lobes. One of the main limitations of segmentation methods that heavily rely on intensity values is the inability to correctly classify the same class having different local distributions, even if inhomogeneity correction methods are applied as pre-processing. To increase our model invariance, we introduce, as an additional data augmentation strategy, a synthetic inhomogeneous field noise. We start by pre-computing a 3D multivariate normal distribution, with zero-means and twice the dimension (for each axis, i.e., 8× the volume) of the original volume. For each training volume, we randomly sample from the 3D multivariate normal distribution a noise volume as big as the former volume. The so-generated noise volume is then summed to the anatomical MRI, adding further variability to the volume intensities and simulating distortions along different directions. In Figure 6, a sketch of the method, when applied on a 2D slice example, is shown (see Supplementary Material for further examples on 1D and 3D cases).

**Figure 6.**
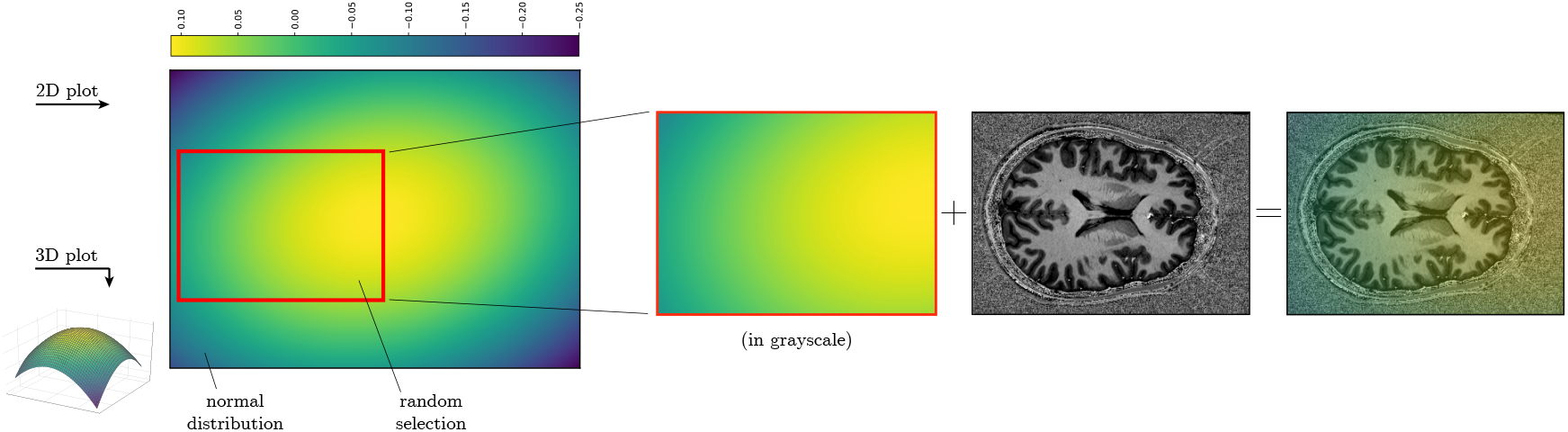
Cartoon to describe the inhomogeneous-field noise for the 2D case. After pre-computing a multivariate normal distribution, we randomly extract a noise sample as big as the sample to augment, for every training batch. The extracted patch is then summed to the original sample.

### 2.3 Manual segmentation

A portion of the considered dataset has been manually annotated by one of the author from the University of Glasgow, who accumulated several years of experience in neuroscience and brain MRI segmentation, and reviewed by a radiologist with 20 years of experience. In particular, 3 subject volumes have been randomly selected from the 7T MRI dataset: for each of them, 8 regions of widely interest in neuroscience - i.e., early visual cortex (EVC), high-level visual areas (HVC), motor cortex (MCX), cerebellum (CER), hippocampus (HIP), early auditory cortex (EAC), brainstem (BST), and basal ganglia (BGA) - have been selected and labelled. Since each of the 8 sub-volumes of interest includes 5 adjacent slices of dimension 150 × 150, the manually labelled dataset accounts for a total number of 2.7*M* voxels (150 × 150 × 5 × 8 × 3).

### 2.4 Deep learning model

Similarly to Bontempi et al. (2020), the model architecture is designed to deal with the dimensionality of the training data (i.e., 256 × 352 × 224 voxels) at once. As shown in Figure 1, the model is a deep encoder/decoder network with three layers, with one, two, and three 3D convolutional blocks in the first, second, and third level respectively (*n*_*filters* = 24, 48, 96).

Since the network is fed with a whole volume as an input, each convolutional block (kernel size 3 × 3 × 3), processes the whole brain structure. The full volume helps the model to learn both local and global structures and spatial features (e.g., the absolute and relative positions of different brain components), which are then propagated to subsequent blocks. Dimensionality reduction is achieved using strided convolutions instead of max-pooling, which contributes to learning the best down-sampling strategy. A dimensionality reduction (of factor 4 on each dimension) is computed after the first layer, to explore more abstract spatial features. Eventually, the adoption of both tensor sum and skip connections, instead of concatenation, helps in containing the dimension of the parameter space to ~ 1.2*M*.

Training, which takes ~24 hours, is performed on a multi-GPU machine equipped with 4 GeForceQ® GTX 1080 Ti, on which different parts of the model are distributed^4^. During training, we optimise the categorical cross-entropy function using Adam (Kingma and Ba, 2014) with a learning rate of 5 · 10^−4^, *β*_1_ = 0.9 and *β*_2_ = 0.999, using dropout (*p* = 0.1 on 2^*th*^ and 3^*rd*^ level), and without batch normalisation (Ioffe and Szegedy, 2015), achieving convergence after ~ 23 epochs. The code is written in TensorFlow and Keras.

## 3 Results

To check the validity of `CEREBRUM-7T` as a segmentation tool for brain 7T data, in Section 3.1, we first provide a quantitative assessment of the obtained segmentation, with and without data augmentation, with respect to the inaccurate labelling obtained by other state-of-the-art methods (iGT). Then, in Section 3.2, we present the outcome of the Turing test carried out on a data portion by experienced neuroscientists who were asked to subjectively evaluate the best segmentations among `CEREBRUM-7T`, the iGT, and a manual segmentation. In Section 3.3 the manual segmentation masks are used as gold standard for the numerical comparison among `CEREBRUM-7T` outcome, the iGT, both FreeSurfer v6 and v7 masks, and the methods from Fracasso et al. (2016) and Huntenburg et al. (2018).

### 3.1 Numerical comparison against iGT

In order to evaluate the effect of the data augmentation strategy, `CEREBRUM-7T` architecture is compared in two variants, with and without data augmentation, against the inaccurate ground-truth. The two models are trained by minimising the same loss and using the same learning rate.

Performance is assessed by three metrics adopted by the MICCAI MRBrainS18 challenge, which are among the most popular ones used in the context of segmentation (Taha and Hanbury, 2015): the Dice Coefficient (DC), a similarity measure which accounts for the overlap between segmentation masks; the Hausdorff Distance computed on its 95th percentile (HD95), which evaluates the mutual proximity between segmentation contours (Huttenlocher et al., 1993); the Volumetric Similarity (VS) as in (Cárdenes et al., 2009), a non-overlap-based metric which considers the similarity between volumes.

The quantitative comparison is outlined in Figure 7 where average results for DC, HD95, and VS) obtained on the 18 test volumes are shown class-wise (i.e., on GM, WM, ventricles, cerebellum, brainstem, and basal ganglia).

**Figure 7.**
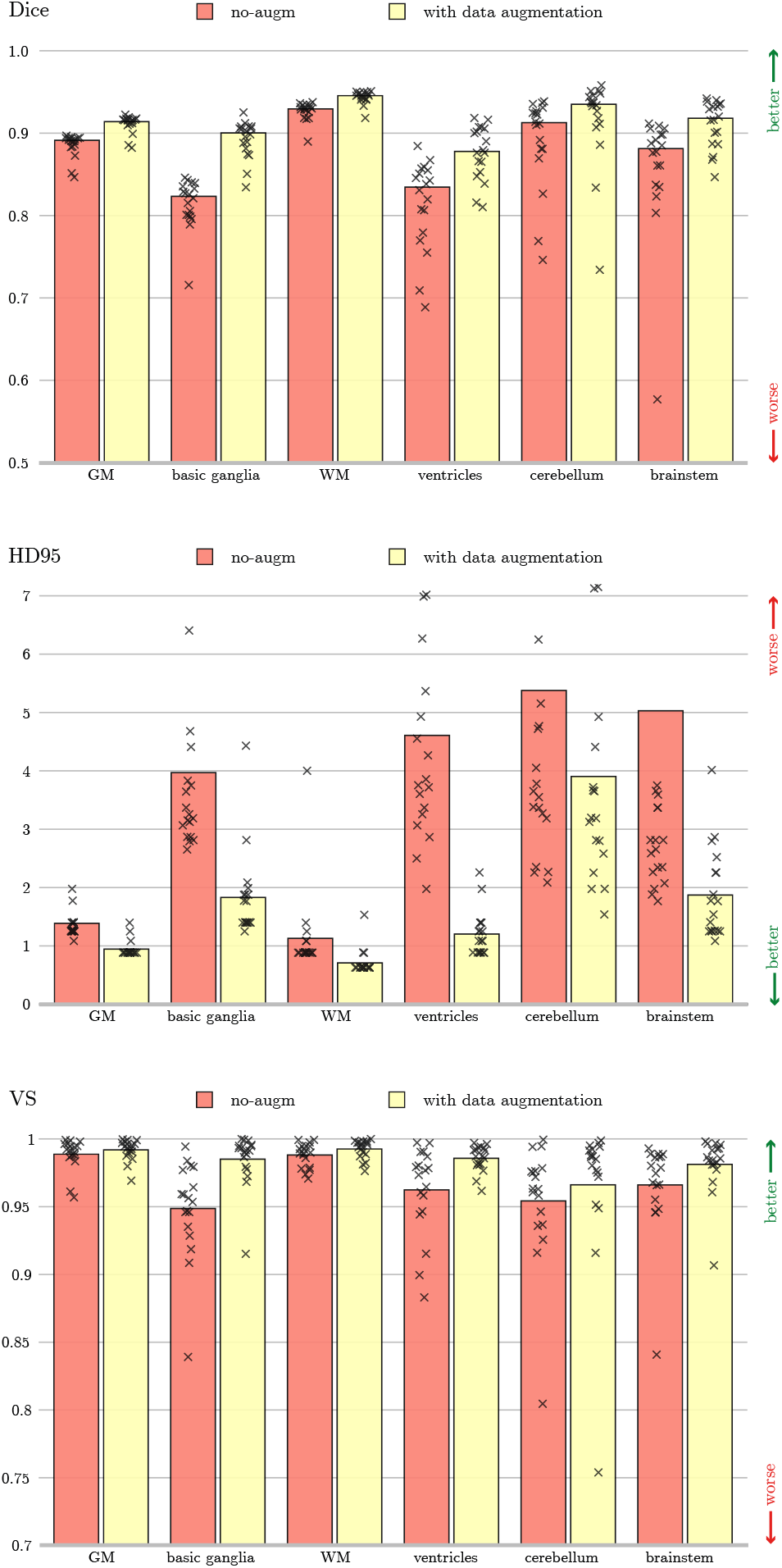
Dice Coefficient, 95th percentile Hausdorff Distance, and Volumetric Similarity computed using the iGT segmentation as a reference. The data augmented model (yellow), and the model trained without data augmentation (red) are compared. The height of the bar indicates the mean across all the test subjects, while every mark is a tested volume.

Overall, the architecture with data augmentation outperforms the baseline solution on every class, independently from the observed metric. This is especially noticeable for HD95, where the difference in the average score between the two configurations (computed on all test volumes) is proportionally more prominent than for other metrics. This might be due either to a larger variability (which might affect the reliability of the measure), or to the fact that, since HD95 accounts for differences in segmentation contours, the beneficial effects given by offline data augmentation (i.e, shifts, rotations, and morphing) reflects on an increased accuracy of the segmentation borders. Such interpretation is supported by the observation that smaller brain structures, such as ventricles, brainstem, etc., where the identification of segmentation boundaries is most critical, are the ones that benefit the most from such augmentation. In summary, this result points out how much the applied data augmentation strategy helps segmentation, significantly improving results from Bontempi et al. (2020).

### 3.2 Turing test: results

From the quantitative assessment presented in Section 3.1 emerges that there is a relative difference in performance between `CEREBRUM-7T` architectures and the iGT used for training. Nevertheless, since performance are measured with respect to an inaccurate ground-truth, `CEREBRUM-7T` segmentations might be even superior to those provided by the inaccurate labelling, as in Bontempi et al. (2020) and Roy et al. (2019), as we also suggest in Figure 2 where `CEREBRUM-7T` masks appear more accurate than iGT masks.

To test this hypothesis, we design a Turing test in which seven expert neuroscientists (different from those who generated the manual segmentation) are asked to choose the most accurate results among three provided ones: the mask produced by `CEREBRUM-7T`, the iGT, and the manual segmentation (intended as gold standard).

If systematically proven, the superiority of `CEREBRUM-7T` against the iGT would confirm the validity of the weakly supervised learning approach, resulting in a learnt model with generalisation capability over its training set obtained with state-of-the-art methods. Furthermore, a human expert evaluation, compared to a purely numerical measure, has the advantage to account for the grade of severity of every single segmentation error, giving important feedback on the suitability of the segmentation for the application (Taha and Hanbury, 2015).

The survey participants are presented with a set of randomly arranged slices taken from the manually annotated dataset: they are either axial, sagittal, or coronal views from the eight selected areas of interests (see Section 2.3 for details) segmented with the three compared methods (`CEREBRUM-7T`, iGT, and manual segmentation). For each presented couple of segmentation results, the expert is asked to choose the best one between the two, or to skip to the next slice set if unsure. Each participant inspects all eight areas of interest, for each of the three test volumes. To better compare results in a volumetric fashion, it is also possible for the participant to browse among neighbouring slices (two slices before, and two after), and interactively change the mask opacity to more easily check for an exact anatomical overlap. A snapshot of the survey interface, coded with PsychoPy (Peirce et al., 2019), is provided in the Supplementary Material.

As evident when inspecting the aggregated results of the Turing test shown in Figure 8(a), the direct comparison between `CEREBRUM-7T` and the iGT shows that survey participants clearly favoured our proposed solution. Moreover, when both `CEREBRUM-7T` and the iGT are compared against manual segmentation, `CEREBRUM-7T` obtains more favourable results than iGT. This is confirmed also when results are split per different brain areas: in the comparison against manual, the iGT is almost never chosen, while in selected areas (i.e., EVC, MCX, HIP) `CEREBRUM-7T` becomes competitive also against the gold standard offered by manual segmentation.

**Figure 8.**
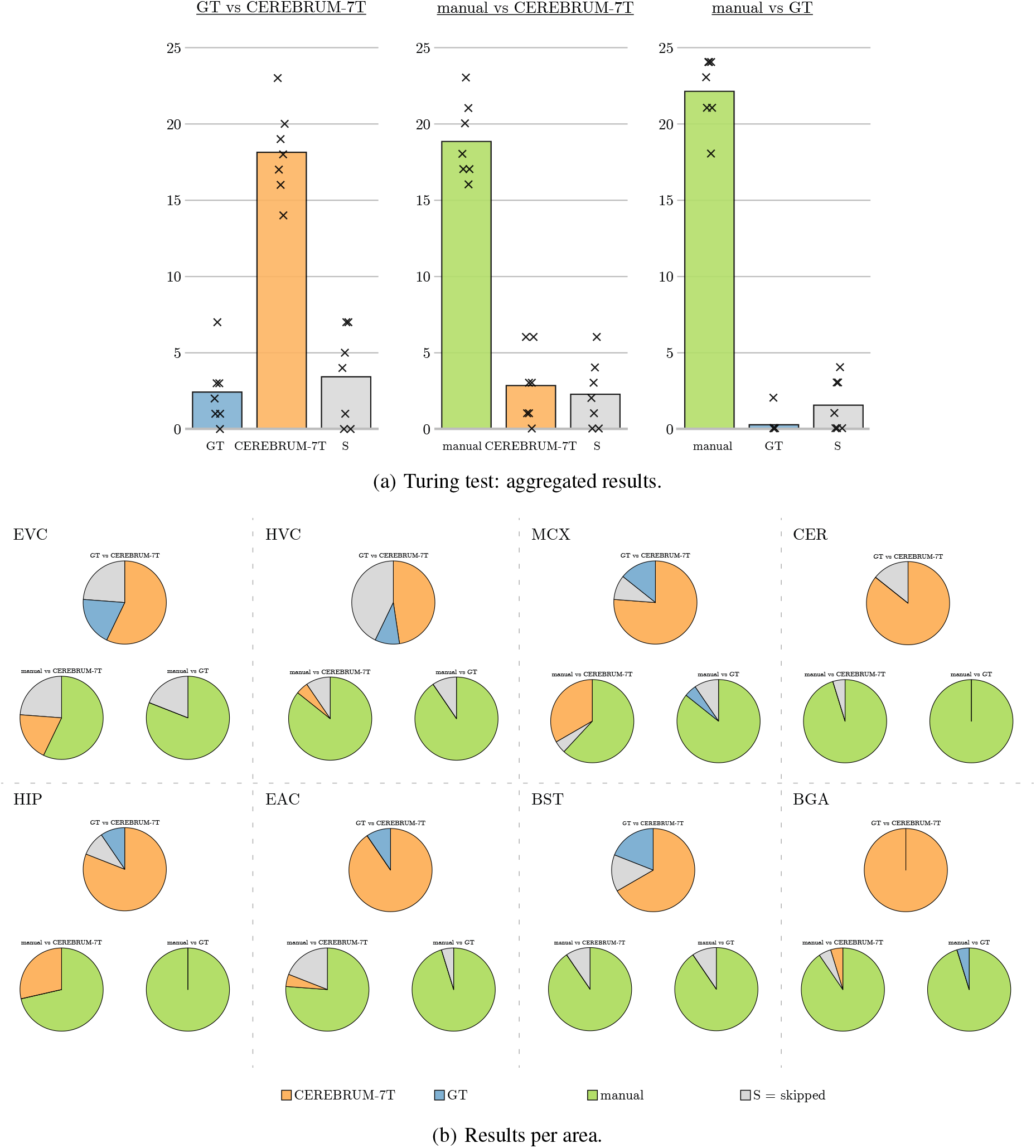
Results of the Turing test. (a) The three subplots show the three comparisons questioned during the survey (manual vs `CEREBRUM-7T`, manual vs iGT, iGT vs `CEREBRUM-7T`), since segmentations masks were presented in couples. iGT votes are displayed in blue, `CEREBRUM-7T` in orange, while skipped responses (S), meaning participants could not choose between the two segmentations, are displayed in grey. The height of bars indicate the means across subjects (i.e, how many times a selection was made, where max. is 3 volumes × 8 areas = 24); every mark x is a participant. (b) Results are split per area of interest: early visual cortex (EVC), high-level visual areas (HVC), motor cortex (MCX), cerebellum (CER), hippocampus (HIP), early auditory cortex (EAC), brainstem (BST), and basal ganglia (BGA).

### 3.3 Numerical comparison against manual segmentation

For the sake of completeness, we present a purely numerical evaluation based on Dice coefficient of the same dataset of 2.7*M* voxels used in the Turing test (three volumes, eight selected areas per volume). Considering manual annotations as a reference, Figure 9 presents a quantitative comparison among `CEREBRUM-7T`, the iGT, the results obtained by FreeSurfer v6 and Freesurfer v7, those by the method presented in Fracasso et al. (2016) (on applicable classes only, i.e., GM and WM), and those from Huntenburg et al. (2018). The results, separately presented for each of the six brain categories, confirm that `CEREBRUM-7T` returns the most accurate segmentation on all brain structures against all other state-of-the-art methods.

**Figure 9.**
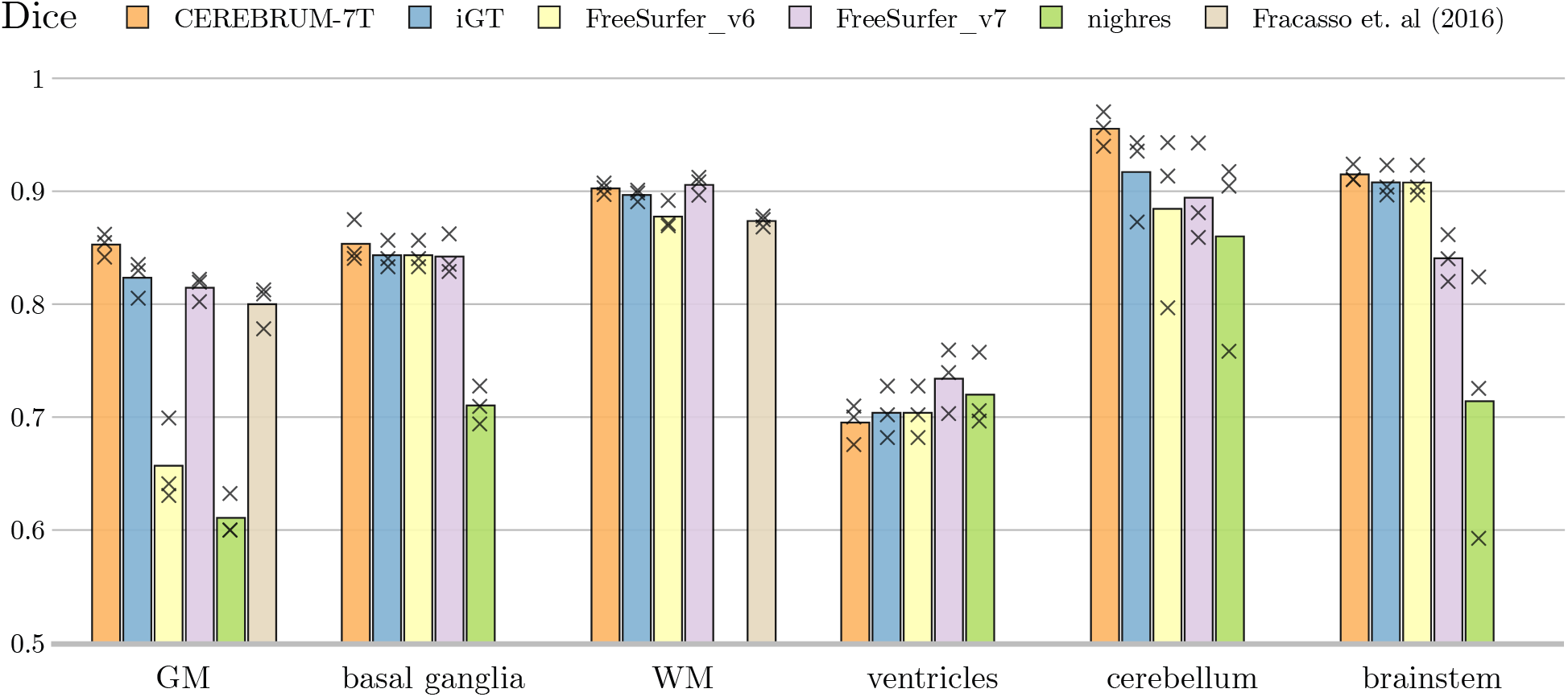
Using manual annotation as a reference, comparison of Dice coefficient between our method (`CEREBRUM-7T`), the iGT used for training, FreeSurfer v6, FreeSurfer v7, and the segmentation tools in Fracasso et al. (2016) (only GM/WM) and Nighres by Huntenburg et al. (2018). Every mark is a tested volume from the manually annotated testing set. Nighres result (green bar) is missing for WM since the Dice coefficient is below 0.5.

## 4 Fine-tuning on data from different sites

We here present a fine-tuning procedure which enables to extend the previously trained model on new datasets, also collected in different sites. It is in fact well known the issue related to the *distribution shift* (Quionero-Candela et al., 2009): different data acquired from different sites usually significantly differ in statistical distribution. A researcher eager to try the model on another independent dataset - which is the usual research setting scientists find themselves in - would face the fact that the DL model would start to struggle, as the data statistics they learnt are not sufficient anymore to carry out the task such models were trained for. For this reason, it is important to fine-tune the model for the new dataset, adapting it to the new statistical distribution of the data.

By the following fine-tuning procedure we describe how to transfer the ability of the method to two new small datasets, using automatic and manual segmentations as fine-tuning labels. All steps of the procedure are detailed in Figure 10: data preparation (step 1) includes operations of rotation and cropping, and requires the computation of the training labels (either by automatic tools or manually) and the mean/std volumes of the new dataset. Afterwards, geometric data augmentation, for both the anatomical and the labelled volumes, are performed offline (step 2). Step 3 describes the “warming-up” of the model, in which the new layer weights are learnt without compromising the frozen layer features obtained during the training (*lr* = 1 · 10^−5^). Finally, step 4 is responsible for the fine-tuning of the entire network (*lr* = 5 · 10^−4^).

**Figure 10.**
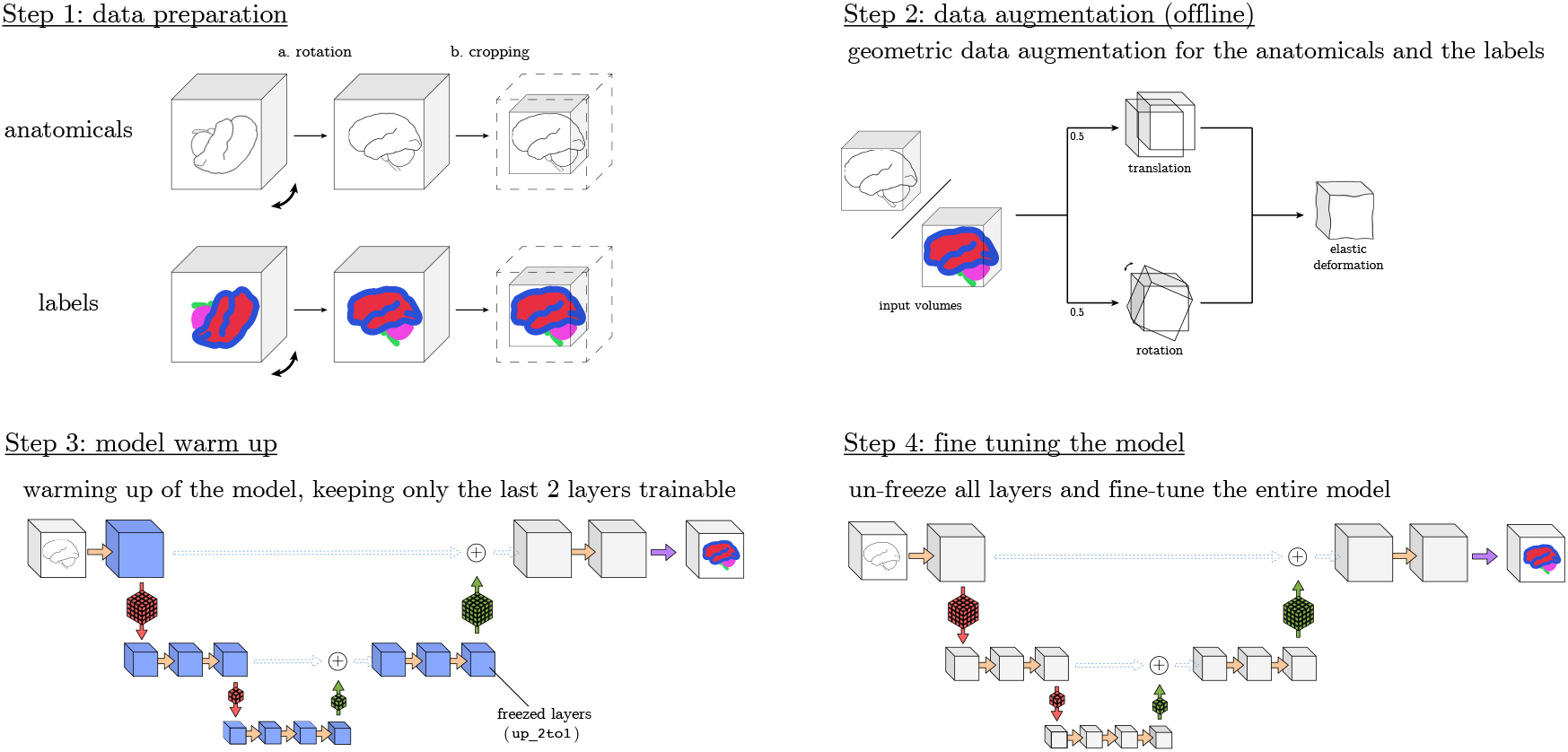
Steps to fine tune the method to a new sequence or site.

### 4.1 Fine-tuning on AHEAD dataset (Alkemade et al., 2020)

We first fine-tune the model trained on Glasgow data by using only 20 volumes from AHEAD dataset (Alkemade et al., 2020). The Amsterdam Ultra-high field adult lifespan database (AHEAD) consists of 105 7T whole-brain MRI scans, including both male and female subjects (age range 18–80 yrs). In order to mimic one real scenario, we use one of the most recent tool openly available on the market: FreeSurfer v07. We select 20 volumes with good segmentation masks, and we augment every volume 15 times. Validation is performed on 4 volumes, while the remaining 81 are used as a testing set. The labels used for training the network are those derived from the iGT, while those used for fine tuning are obtained by FreeSurfer v7 (Fischl, 2012), to completely disentangle the model from the method in Fracasso et al. (2016).

#### 4.1.1 Results on AHEAD data

Although only 20 volumes were used for fine-tuning the model (previously trained on Glasgow data), the improvements in the results with respect to using CEREBRUM-T7 off the shelf are striking, as shown in Figure 11, which presents such comparison (also versus FreeSurfer v7) on slices from 5 different subjects of the AHEAD testing set.

**Figure 11.**
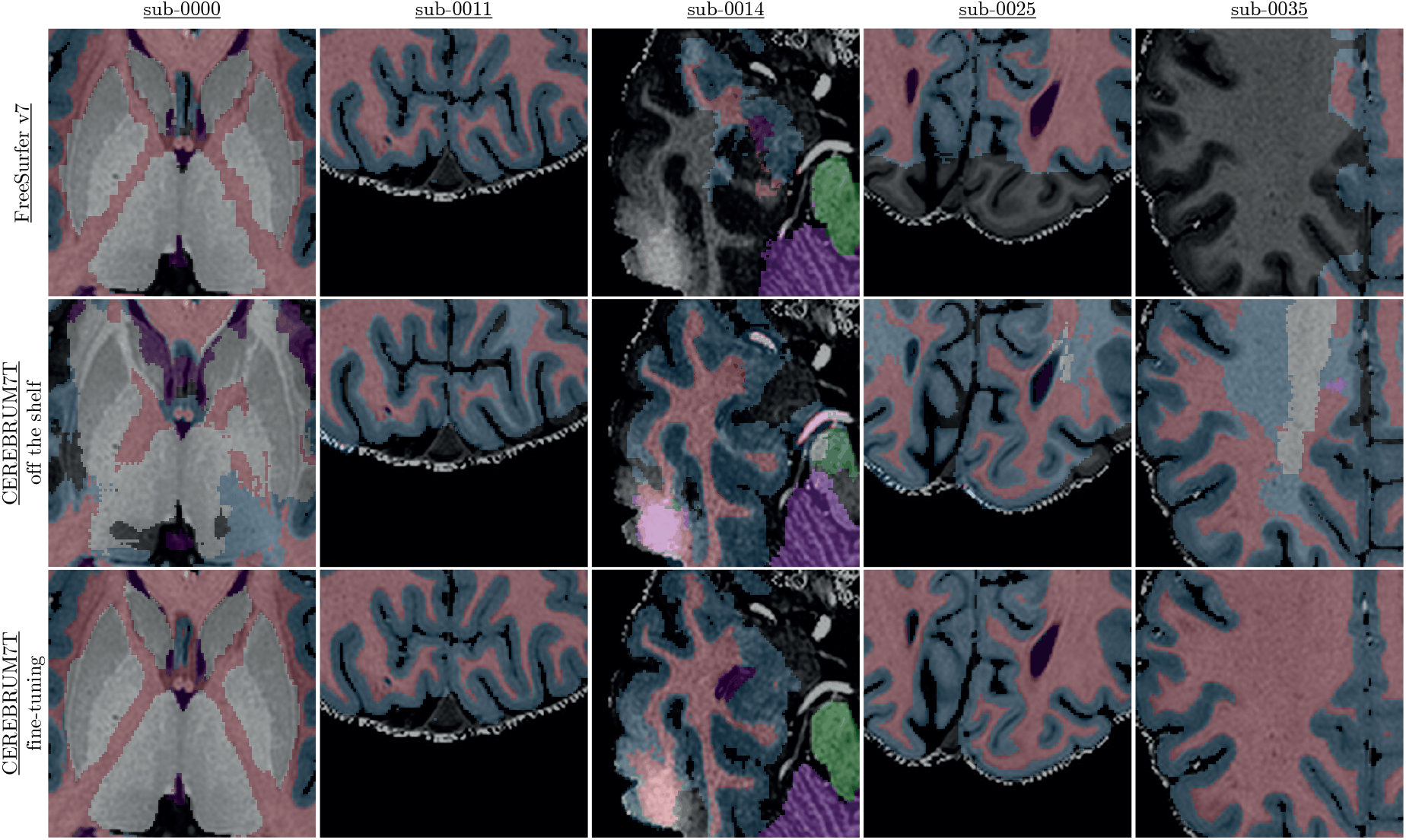
Fine tuning results on AHEAD data: comparison between FreeSurfer v7, plain CEREBRUM-T7 (trained on Glasgow data and tested on AHEAD data), and CEREBRUM-T7 fine-tuned for AHEAD data. More results in Supplementary. Animated GIF on the project website.

In particular, in the comparison versus Freesurfer 7, it is evident that `CEREBRUM-7T` produces much smoother masks. Whereas FreeSurfer v7, which has been improved for UHF data, is able to segment very well multiple areas (e.g., GM/WM boundary), the inhomogeneity of the scan still affects its ability to correctly select all regions (e.g., the parietal and temporal lobes), often producing holes in the segmentation masks (see the project website for more results).

### 4.2 Fine-tuning on Schneider et al. (2019) dataset

In this final experiment, we investigate the limitations of our method deploying it in the following challenging scenario: we fine-tune the model (trained on Glasgow data) to predict the manual segmentation provided in Schneider et al. (2019). Such dataset contains sub-millimetre 7T MRI images of the human brain, which are thought for the supervised training of algorithms to perform tissue class segmentation. In particular it includes preprocessed MRI images (co-registered + bias corrected) and corresponding ground truth labels: white matter, grey matter, cerebrospinal fluid, ventricles, subcortical, vessels, and sagittal sinus. For our purposes we exploit only the MP2RAGE subset, which exhibits excellent manual segmentation masks based on 4 subjects.

Since the dataset contains only 4 volumes, we exploit a cross-validation strategy, using 3 volumes for training and 1 for testing. Since the method needs a larger training set, we apply a strong augmentation procedure, concatenating different volume manipulation strategies: translation, rotation, and morphing. Doing so, we create 30 volumes for each training sample (for a total of 90). Due to technical limitations, we decided to fine-tune the model on six classes only: white matter, grey matter, cerebrospinal fluid, ventricles, subcortical, and vessels. To ease the task, we also apply a brain mask to the volume, cropping outside the skull.

#### 4.2.1 Results on Schneider et al. (2019) data

Results show that, with only 3 volumes for fine-tuning, the predicted labels are very accurate (see Table 2), which shows that the model provides great flexibility. To analyse results, we need to distinguish two different cases. When we consider classes which are already known by the `CEREBRUM-7T` model, such as GM, WM, and ventricles - or a combination of previous classes - such as CSF or subcortical (which is a combination of basal ganglia, brainstem, and cerebellum), the model takes advantage of the previous learning (on Glasgow data) and simply transfers/applies the knowledge on the new dataset, producing very accurate results. Conversely, when inferencing on classes never seen before, like vessels, on which the model has not a prior-knowledge, segmentation results are qualitatively lower, as expected. Visual results can be inspected on the project website.

**Table 2.** Dice coefficient computed on the 4 brain volumes of Schneider et al. (2019) data.

| Subject | sub-001 | sub-013 | sub-014 | sub-019 |
| --- | --- | --- | --- | --- |
| Dice coeff. (tot) | 0.977 | 0.977 | 0.980 | 0.977 |

## 5 Discussion

In this work, we design and test `CEREBRUM-7T`, an optimised end-to-end CNN architecture that allows the segmentation of a whole MRI brain volume acquired at 7T. The speed of computation, i.e., few seconds, and the quality of obtained results, make `CEREBRUM-7T` the most advantageous solution for 7T MRI brain segmentation among the few currently available. Furthermore, as shown above, by following a simple fine-tuning procedure, any research in the field is able to use `CEREBRUM-7T` to segment brain data from different MRI sites. Eventually, in order to allow other researchers to replicate and build upon `CEREBRUM-7T` findings, we make code, 7T data, and other materials (including iGT and Turing test) available to readers.

### Full volumetric segmentation

Similarly to `CEREBRUM`, also `CEREBRUM-7T` processes the whole brain volume as one, avoiding the drawbacks of the tiling process (Reina et al., 2020), thus preserving both global and local contexts. This partially resembles what happens during manual segmentation: first, the expert looks at the brain volume from afar to identify where different brain structures are located (global clues). Once a coarse segmentation is apparent, the expert begins to segment voxel by voxel at the pixel level, focusing only on a specific area (local processing). For a human, both of these two levels (or scales) of information are needed to perform the segmentation.

`CEREBRUM` preserves such two-scale analysis: global features are obtained by analysing the volume at once, without partitioning. The full-resolution processing of the first layer enables to perform a maximum resolution analysis. A table reporting the receptive fields for each convolutional block of `CEREBRUM-7T` can be found in the Supplementary Material (Table 1).

Classical automatic (pre-DL) segmentation tools, instead, emulate these two steps using atlases to gain global clues and, for most of them, gradient methods for the local processing. For what concerns DL segmentation methods based on tiling, they conceptually lack in the gain of global clues. Furthermore, limitations in memory size of accelerator cards, prevented so far large medical volumes from being processed as a whole: thanks to the reduction of network layers we applied on the model architecture, it was possible to make the exploitation of global spatial information computationally tractable. In fact increasing the depth of a CNN does not always allows the model to capture richer structures and yielding better performance. On the contrary, as highlighted by works such as Perone et al. (2018), in some cases the low-level features extracted by the network prove to be the most important ones - even if the task is complex. Maintaining a small number of layers allow us to analyse the volume at full resolution and at once, gaining both global and local scale: this brings in a sense our DL model closer to an atlas, with respect to any other previous approach, since it finally learns a-priori probabilities for every voxel.

### Mesh reconstruction

Our `CEREBRUM-7T` approach is able to efficiently produce 3D high-quality models useful for the neuroscientific and biomedical communities. For example, in functional MRI (fMRI) studies, researchers need first to isolate specific brain structures (e.g., GM) in order to analyse the spatio-temporal patterns of activity happening within it. As such, we show in Figure 12 a view on four reconstructed meshes (WM/GM boundary and outer GM boundary) obtained from a testing volume of the independent dataset AHEAD, processed by FreeSurfer V7 (left) and `CEREBRUM-7T` (right), respectively.

**Figure 12.**
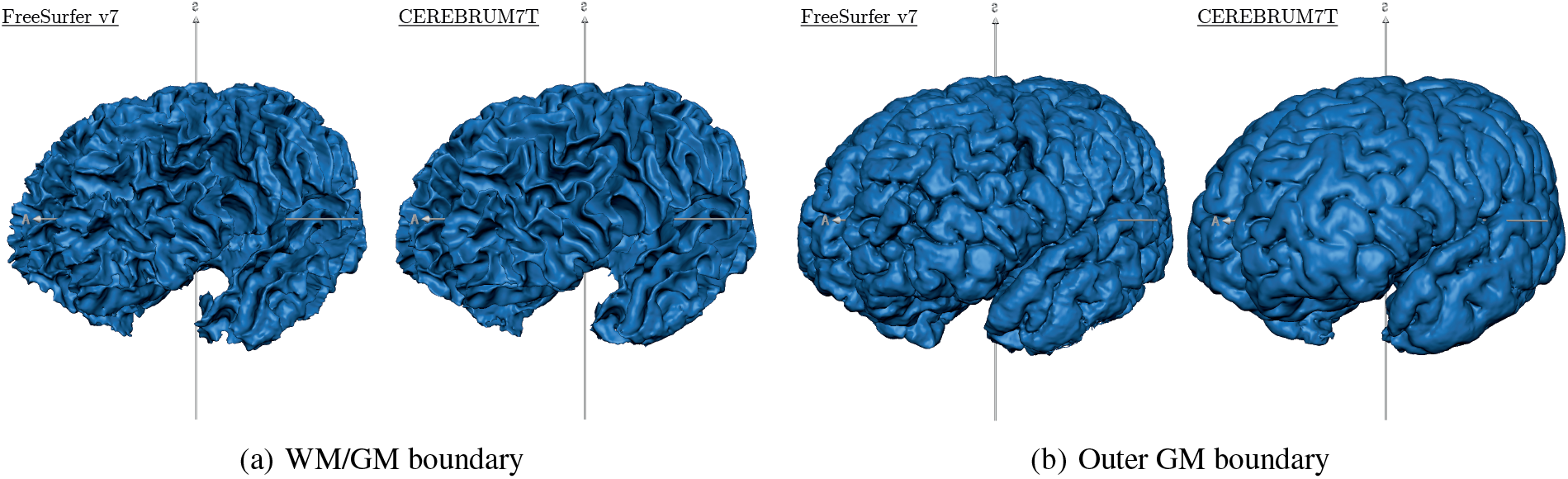
Reconstructed meshes of (a) WM/GM boundary and (b) outer GM boundary of a testing volume of the independent dataset AHEAD - sub. 88 - for FreeSurfer V7 (left) and CEREBRUM (right). A light smoothing operation is performed on both meshes (50 iterations - BrainVoyager, Brain Innovation (Goebel, 2012)) - no manual corrections performed. We added un-smoothed meshes on Supplementary. More results (animated GIF) on the website page.

By operating as a true 3D structure model, `CEREBRUM-7T` ensures globally smoother and more coherent surfaces across slices with respect to 2D methods, both manual and automatic. Commonly adopted editing tools, such as ITK-SNAP (Yushkevich et al., 2006) or `ilastik` (Berg et al., 2019), usually display 3 synchronised 2D orthogonal views onto which the operator draws the contour of the structures. The extraction of a continuous 3D surface from the collection of 2D contours, as well as from 3D tiles, is a nontrivial postprocessing task, where bumps in the reconstructed 3D surface are often inevitable due to inter-slice inconsistencies in segmentation.

### Probability maps

To further appreciate the quality of `CEREBRUM-7T` output, in Figure 13 we show the segmentation inferred by the model before thresholding (percent probability). In such probability maps, each voxel intensity is associated with the probability of belonging to the most likely class. Since most voxels are associated with significant probability of belonging to their correct brain class, such maps demonstrate the ability of the model to make use of both global and local spatial cues. Furthermore, the almost total absence of spurious activations, confirms the high level of confidence achieved by the model. In testing, the model outputs both the probability maps and the thresholded segmentation mask by default.

**Figure 13.**
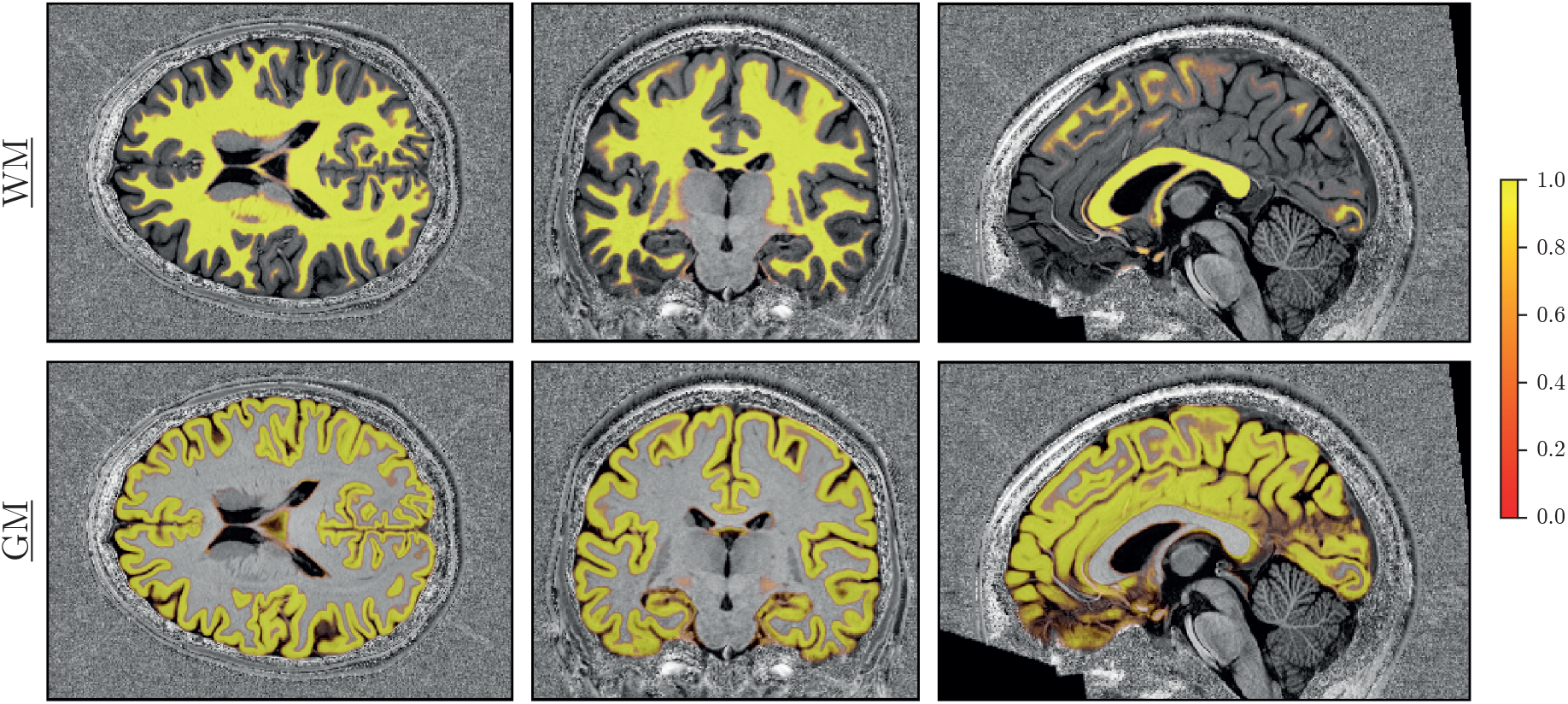
Soft segmentation maps (i.e., probability maps) of a testing volume for WM and GM classes. The model produces maps very consistent probabilities, giving additional flexibility than a hard thresholded map. Remaining classes are shown in Supplementary Material.

### Fine-tuning

Results shown in fine-tuning experiments are notable because they directly tackle one of the main limitation of deep learning: the need for big datasets. Although it is pretty straightforward to say that, in such scenario (i.e., very limitated training samples), safer strategies can be applied, - like decomposing the volumes in slices and apply a slice-based method -, we still demonstrate the practical portability of the trained model and a procedure compliant with the objective of releasing a tool which is effective and efficient also on data from new sites.

### Limitations

Having decided to process the whole volume at once, which required to maintain a model with low level of complexity, it was not possible to include network elements which are very popular in recent deep learning architectures (e.g., dense layers). As another downside of the choice of processing the whole brain volume at once, it was not possible to increase the batch size to a value greater than one, due to the technological constraints of GPU memory. We chose to analyse the volume in its entirety, instead of exploiting the advantages that the increase of the batch size could carry. With the rapid increase of hardware capabilities, we are confident to be able soon to manage more recent architecture elements, and larger batch sizes.

Being developed on proprietary data, and especially inserted in a study pipeline mostly focused on WM/GM analysis, we performed neck cropping on data. Such reduction in the size of the input volume allowed for an increase in the filter number, increasing model capability. To provide another example of this, additionally showing the flexibility of the method, we crop both T1_w_ and iGT on the visual cortex and we retrain the model on GM and WM classes only. Segmentation results are presented in the Supplementary Material.

Last, our choice to perform segmentation only on one sequence (i.e., T1_w_) was made in order to limit the scanning time, which is a constraint often imposed for reducing the patient discomfort. This choice also avoids the need for sequence alignment, and the reduction of distortion and morphing which are typical of each sequence. However, for whoever would like to develop a segmentation method combining multiple sequences, the code provided can be easily extended to other sequences without adding much more complexity to the model.

## Acknowledgements

This project has received funding from the European Union Horizon 2020 Framework Programme for Research and Innovation under the Specific Grant Agreement No. 785907 and 945539 (Human Brain Project SGA2 and SGA3).

We thank Dr. Alessio Fracasso for assistance with the inaccurate ground-truth creation, the Muckli lab for the Turing test and neuroimaging knowledge, Dr. Mattia Savardi for very useful technical comments, Dr. Alberto Signoroni for fruitful discussion on learning strategies, Dr. Lucy Petro for comments improving the manuscript, and Mrs Frances Crabbe for useful feedbacks on manual segmentation.

## Author Contributions

**Michele Svanera**: Conceptualisation, Methodology, Software, Validation, Formal Analysis, Investigation, Data Curation, Writing - Review & Editing, Visualisation. **Sergio Benini**: Conceptualisation, Writing - Original Draft, Writing - Review & Editing, Supervision, Project Administration. **Dennis Bontempi**: Conceptualisation, Methodology, Software, Writing - Original Draft, Writing - Review & Editing. **Lars Muckli**: Resources, Funding Acquisition.

## Footnotes

2 With the term “out-of-scanner” we refer to the reconstructed data saved in DICOM 2D images.

3 Data are openly available under EBRAINS knowledge graph.

4 Other graphic cards are also suitable for the purpose, such as 1×RTX 8000 or 2×RTX 3090.

