## Supplementary Material for "CEREBRUM-7T: Fast and Fully-volumetric Brain Segmentation of 7 Tesla MR Volumes"

A Preprint

December 3, 2020

#### 1 Survey programming

As we discussed in Section *3.2. - Survey results* of the manuscript, we designed and implemented a PsychoPy (Peirce et al., 2019) test for neuroscientists to assess the segmentation quality of CEREBRUM-7T comparing our model with manual segmentation and with the GT. In Figure 1, we show the interface we presented the participants with. The source code to replicate the test is made available at the Project GitHub page[1].

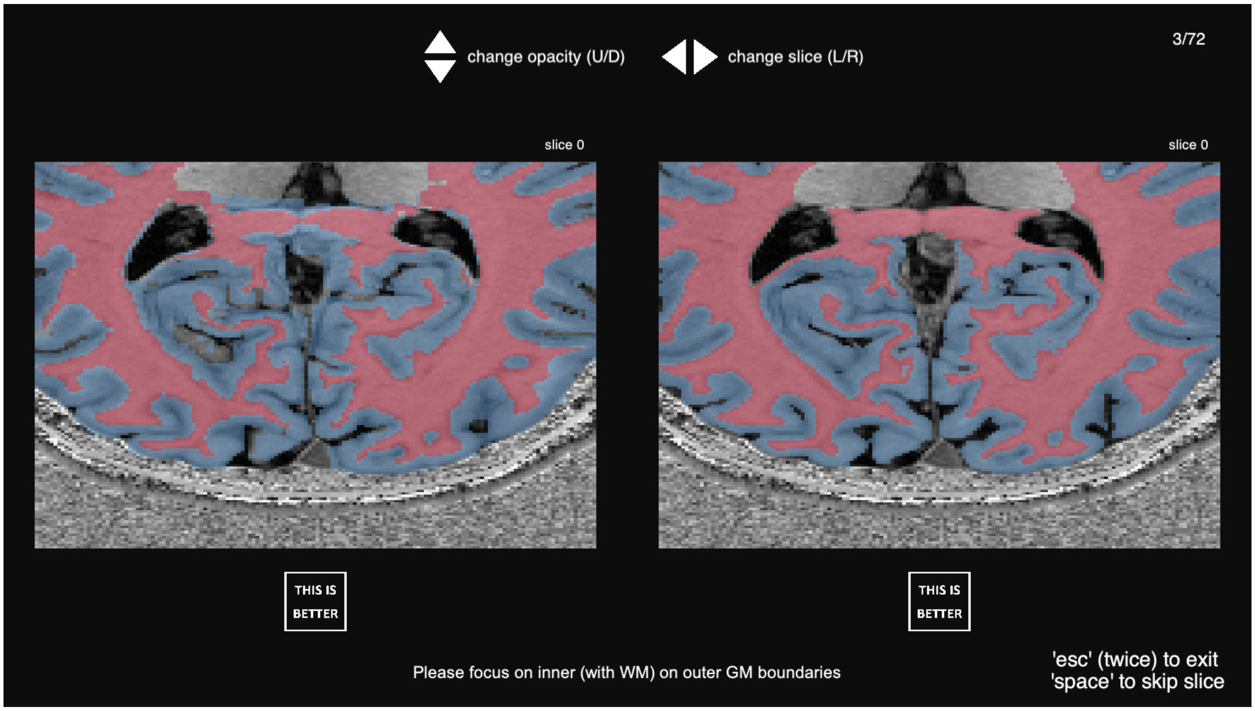


Figure 1. Survey interface. Two different segmentations of the same anatomical section are displayed. The participant needs to focus on specific classes (declared by the instructions on the bottom) and select the best segmentation (or skip). The participant has also the ability to change the opacity of the segmentations with up/down arrows and look at previous and next slices with left/right arrows.

[1] https://github.com/rockNroll87q/cerebrum7t

#### 2 Network receptive field

In Table 1 we report the receptive field for every convolutional block of `CEREBRUM-7t`. The capability of the model to learn both local and global features is coherent with the last layer units of the model having a $100 \times 100 \times 100$ theoretical receptive field.

Table 1
Architecture details and theoretical receptive field. All the convolutional blocks exploit a $3 \times 3 \times 3$ kernel as a feature extractor, unless otherwise specified.

| Block | Input Size | Output Size | Receptive Field | Channels (In/Out) |
|---|---|---|---|---|
| L1 - conv3D - #1 | $256 \times 352 \times 224$ | $256 \times 352 \times 224$ | $3 \times 3 \times 3$ | $1/24$ |
| L1 - stridedConv | $256 \times 352 \times 224$ | $64 \times 88 \times 56$ | $6 \times 6 \times 6$ | $24/48$ |
| L2 - conv3D - #1 | $64 \times 88 \times 56$ | $64 \times 88 \times 56$ | $14 \times 14 \times 14$ | $48/48$ |
| L2 - conv3D - #2 | $64 \times 88 \times 56$ | $64 \times 88 \times 56$ | $22 \times 22 \times 22$ | $48/48$ |
| L2 - stridedConv | $64 \times 88 \times 56$ | $32 \times 44 \times 28$ | $26 \times 26 \times 26$ | $48/96$ |
| L3 - conv3D - #1 | $32 \times 44 \times 28$ | $32 \times 44 \times 28$ | $42 \times 42 \times 42$ | $96/96$ |
| L3 - conv3D - #2 | $32 \times 44 \times 28$ | $32 \times 44 \times 28$ | $58 \times 58 \times 58$ | $96/96$ |
| L3 - conv3D - #3 | $32 \times 44 \times 28$ | $32 \times 44 \times 28$ | $74 \times 74 \times 74$ | $96/96$ |
| L3 - Conv3DTranspose | $32 \times 44 \times 28$ | $64 \times 88 \times 56$ | $78 \times 78 \times 78$ | $96/48$ |
| L2 - conv3D - #3 | $64 \times 88 \times 56$ | $64 \times 88 \times 56$ | $86 \times 86 \times 86$ | $48/48$ |
| L2 - conv3D - #4 | $64 \times 88 \times 56$ | $64 \times 88 \times 56$ | $94 \times 94 \times 94$ | $48/48$ |
| L2 - Conv3DTranspose | $64 \times 88 \times 56$ | $256 \times 352 \times 224$ | $97 \times 97 \times 97$ | $48/24$ |
| L1 - conv3D - #2 | $256 \times 352 \times 224$ | $256 \times 352 \times 224$ | $100 \times 100 \times 100$ | $24/24$ |
| L1 - conv3D - #3* | $256 \times 352 \times 224$ | $256 \times 352 \times 224$ | $100 \times 100 \times 100$ | $24/7$ |

* convolutional block preceding the softmax layer, $1 \times 1 \times 1$ kernel.

#### 3 Inhomogeneous-field noise creation

In Figure 2, a sketch of the method for cases 1D, 2D, and 3D. In the manuscript, at Figure 6, only the case 2D is reported.

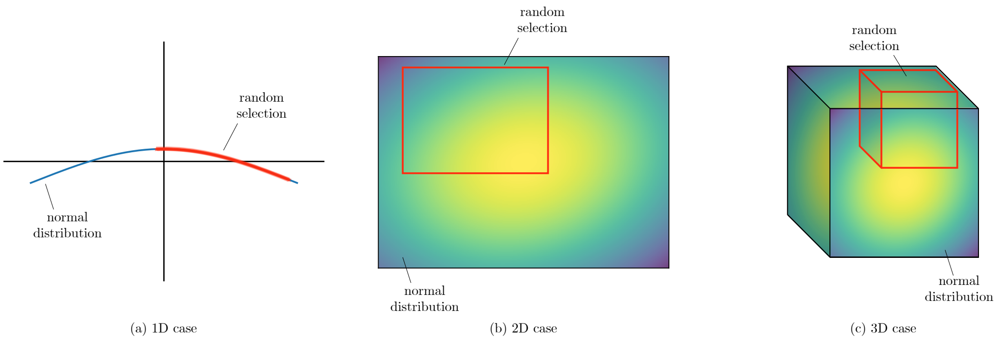


Figure 2. Cartoon describing the inhomogeneous-field noise for the 1D, 2D, and 3D cases.

### 4 Survey results

In Figure 3 we show the results of the survey with participants' details.

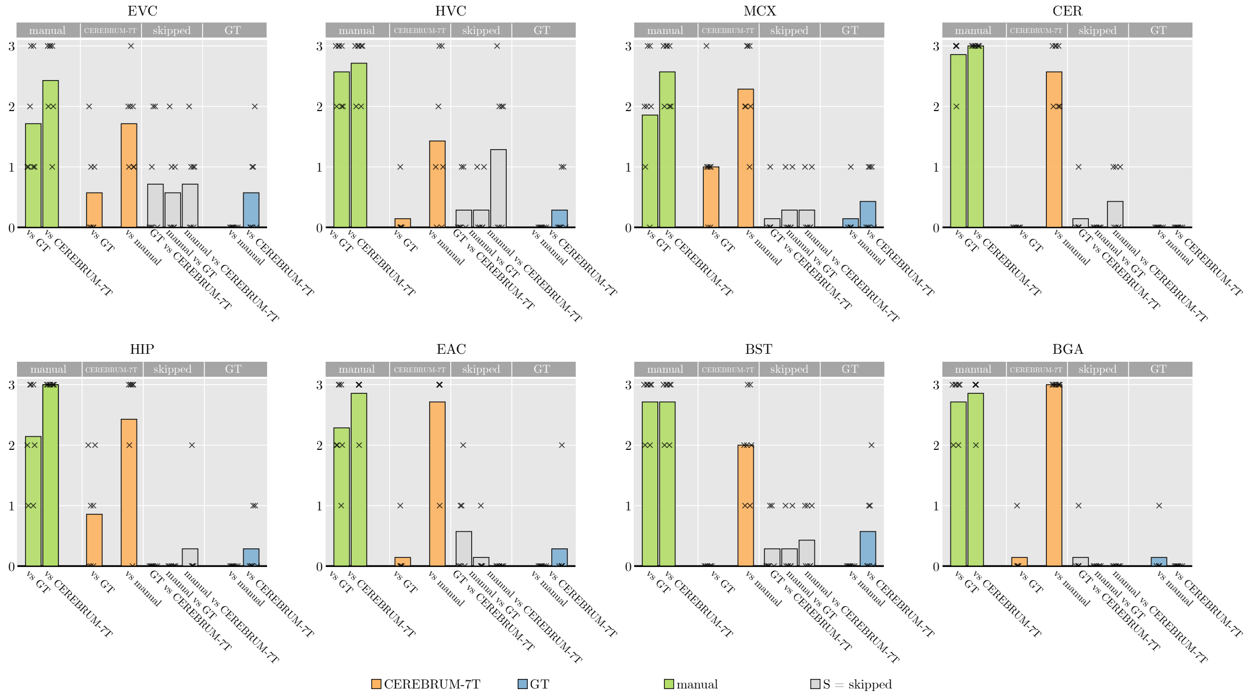


Figure 3. Results of the survey. Results are decomposed per area of interest: early visual cortex (EVC), high-level visual areas (HVC), motor cortex (MCX), cerebellum (CER), hippocampus (HIP), early auditory cortex (EAC), brainstem (BST), and basal ganglia (BGA). Expanding Figure 8-b of the manuscript with detail of participants' agreements.

### 5 Segmentation on a specific area of interest

As discussed in manuscript *Discussions*, subsection *Future extensions and applications*, in some functional studies, it is important to have a model capable to segment only a specific area (for example hippocampus), since, in order to save scanning time, the anatomical scan is sometimes acquired only on the area of interest. To provide an example of the flexibility of the method, we crop both $T1_w$ and GT on the visual cortex and we retrain the model on GM and WM classes only. Results are shown in Figure 4.

### 6 Mean and std volumes for z-scoring

In Figure 5, the z-scoring volumes of denoised data are shown.

### 7 Probability maps

In Figure 6, we show the probabilities maps for the remaining labels: ventricles, basal ganglia, brain stem, cerebellum, and background as well. The figure extends the Figure 11 of the manuscript.

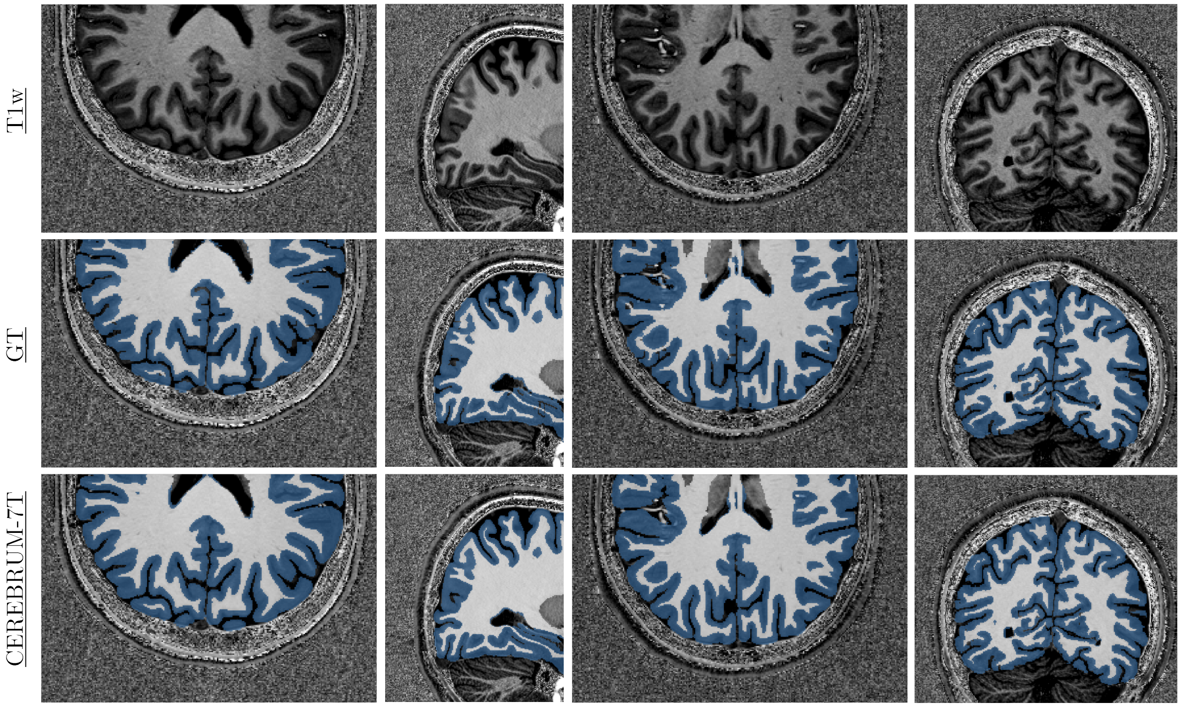


Figure 4. Segmentation of visual cortex for WM and GM classes only. Every column is a different testing volume. Rows are T1w, GT, and the method prediction output.

#### 8 Probability maps

We show in Figure 7 a view on reconstructed meshes for GM and WM obtained from a testing volume processed by `CEREBRUM-7T`. The meshes are reconstructed by using Brainvoyager (Goebel, 2012) directly from both the inner and outer GM boundaries, without any manual correction; the right one was subjected to a light smoothing operation performed on 50 iterations to enhance the visual quality of the 3D model.

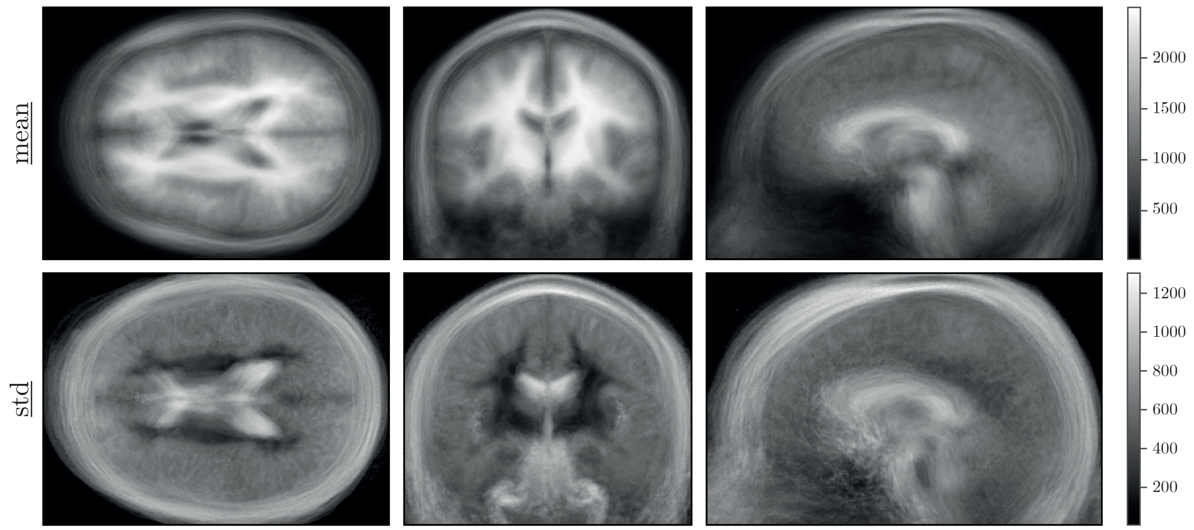


Figure 5. Mean and standard deviation volumes of the denoised database. The original z-scored volumes (not denoised) are in the manuscript in Figure 4.

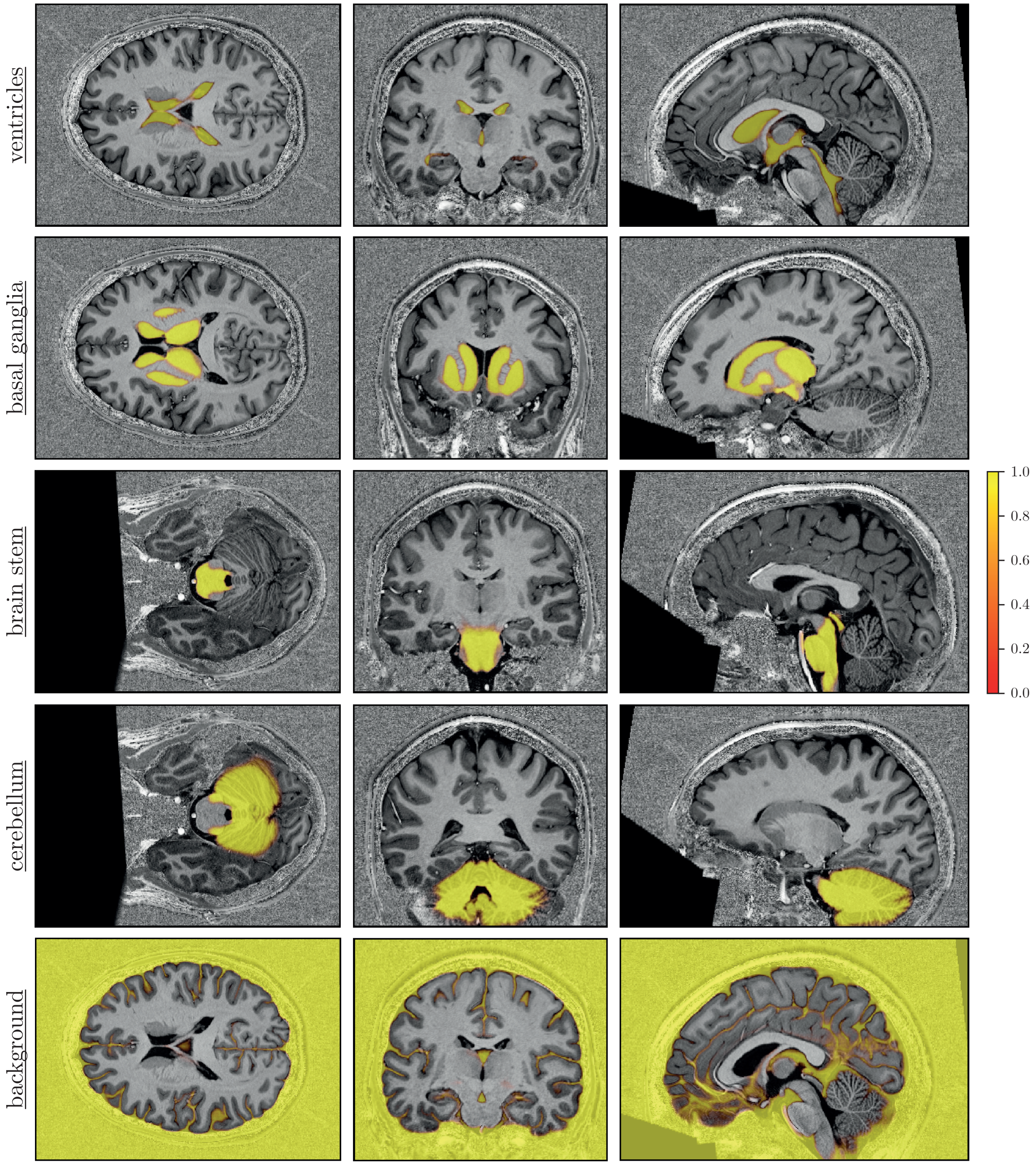


Figure 6. Soft segmentation maps (i.e., probability maps) of a testing volume for classes: ventricles, basal ganglia, brain stem, cerebellum, and background.

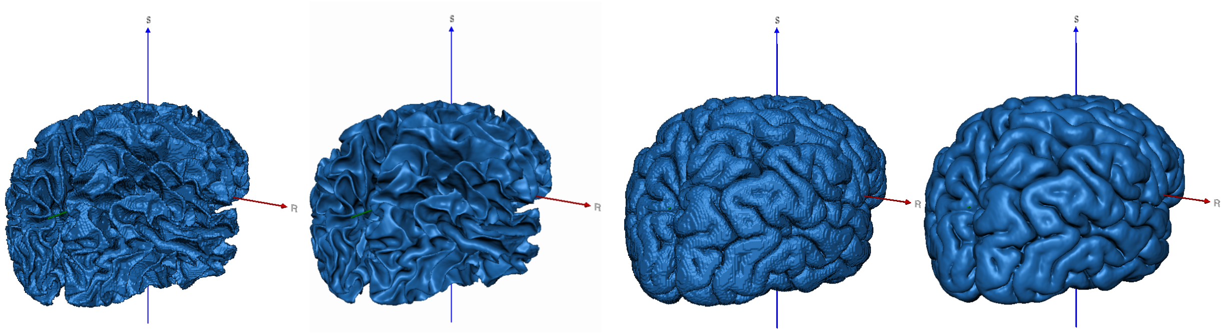

(a) WM/GM boundary

(b) Outer GM boundary

Figure 7. Reconstructed meshes of (a) WM/GM boundary and (b) outer GM boundary of a testing volume (BrainVoyager, Brain Innovation, Maastricht, The Netherlands). A light smoothing operation is performed on the right mesh (50 iterations) - no manual corrections performed. More results on the website page.
